# Sall1a, a transcription factor responsible for the Townes-Brocks syndrome, is involved in hair cell maturation

**DOI:** 10.64898/2026.11.20.768075

**Authors:** Nerea Montedeoca, Francesca Manocchio, Laura Fargas, Berta Alsina

## Abstract

Hair cells of the inner ear are highly specialized mechanosensory cells that convert auditory and vestibular stimuli into electrical signals essential for hearing and balance. While early hair cells fate commitment is well defined, the transcriptional regulators acting at later stages of hair cell maturation remain poorly understood. Spal-like (SALL) family transcription factors are evolutionarily conserved and implicated in multiple developmental processes, and mutation in humans causes Townes-Brocks syndrome, associated with hearing impairment. However, the role of Sall1 in inner ear development has little been investigated. We demonstrate that *sall1a* is specifically expressed in hair cells after the expression of the early differentiation transcription factors, Pou4f3 and Gfi1. We generated a CRISPR Cas9 knockout of *sall1a* gene in zebrafish, finding that s*all1a* is dispensable for hair cell specification but is essential for hair cell maturation and function. Sall1a deficient larvae display a marked reduction in active hair cells, severe balance defects and impaired startle response. Transcriptomic analysis of hair cells reveals that Sall1a promotes the expression of genes associated with mature hair cell identity and function. Our findings identify Sall1a as a key transcriptional regulator and provide insights into the auditory and vestibular dysfunction observed in Townes-Brocks syndrome.

## 1. Introduction

Townes-Brocks syndrome (TBS) is a rare autosomal-dominant disorder characterized by dysplastic ears, imperforate anus, and thumb malformations, with additional manifestations including renal hypoplasia, congenital heart malformations and high-frequency sensorineural hearing loss, which occurs in approximately 62% of affected individuals (Kohlhase et al., 1999; Leduc et al., 2025). In humans, TBS is caused by mutations in *SALL1* (SAL-like 1 gene) which encodes highly evolutionarily conserved transcription factors belonging to the Spalt-like (SALL) protein family. All SALL proteins have four conserved C2H2 zinc finger binding domains and a N-terminal domain that mediates repression via the binding of chromatin remodelling factors (Álvarez et al., 2021; De Celis and Barrio, 2009; Netzer et al., 2001). Over 60 mutations in SALL1 associated with TBS have been identified (Miller et al., 2012), with most occurring in the “Hotspot” region between the glutamine-rich domain and the first C2H2 ZNF pair (Botzenhart et al., 2007). Most TBS-associated mutations produce truncated proteins that can still interact with other SALL proteins via the glutamine-rich domain, potentially blocking their functions through a dominant-negative mechanism. However, some mutations result in truncated proteins that lack the glutamine-rich domain but retain the C2HC zinc finger motif and the N-terminal repressor domain, indicating that the glutamine domain is not essential for typical TBS symptoms (Bozal-Basterra et al., 2018). This hypothesis is supported by studies in mice, where expression of a truncated SALL1 protein lacking all double zinc fingers but preserving the N-terminal region recapitulates the abnormalities seen in human TBS. For instance, heterozygous *Sall1*-deficient mice display high-frequency sensorineural hearing loss, underscoring the role of *Sall1* in inner ear development and reinforcing the hypothesis that TBS can also result from the dominant-negative effect of truncated SALL1 proteins (McLeskey Kiefer et al., 2003).

However, patients with heterozygous deletions of the entire *SALL1* exhibit a milder form of TBS, which is notable given that homozygous *SALL1* mutations are embryonically lethal in humans. This suggests that haploinsufficiency alone is insufficient to cause the full range of severe, classical symptoms (Borozdin et al., 2006). Despite the well-established hearing deficit in TBS patients, the precise role of SALL1 in the inner ear development remains unknown.

Auditory and vestibular mechanical stimuli are transformed into electrical signals by hair cells (HCs) located in the inner ear, which are then propagated by otic neurons to the brainstem (Gillespie and Müller, 2009). HCs are highly specialized cells that contain a high number of distinct ion channels, a highly organized stereocilia at their apical surface with mechanoelectrical transducer receptors. Specific cadherins and myosins, maintain stereocilia structure and coordinate with the mechanotransduction machinery (Krey and Barr-Gillespie, 2019; McGrath et al., 2017).

During embryogenesis, HCs arise from Sox2^+^ sensory progenitors located in the sensory domains of the inner ear epithelium. The basic helix-loop-helix transcription factor Atoh1 initiates the commitment program to HCs by activating other transcription factors, including Pou4f3, which in turn activates Gfi1 in vertebrates (Hertzano et al., 2004). However, other downstream transcription factors involved in HCs maturation remain poorly defined. In the mouse organ of Corti, Sall1 emerged as a Fgf20-downstream target, its disruption reduced the number of outer hair cells (Yang et al., 2020). In the present study, we show the expression of *sall1a* in differentiating HCs of the zebrafish inner ear after the expression of *pou4f3* and *gfi1.1*. To further disclose its function in HCs, we generated a Cas9-mediated stable knock-out of *sall1a*. Sall1a absence caused a dramatic reduction in the number of active HCs. Moreover, larval behaviour was compromised, and *sall1a* KO larvae did not maintain a standing position and displayed a defective startling response. Transcriptomic analysis of HCs in mutant embryos compared with wild-type (WT) embryos revealed direct target genes of *sall1a* in HCs. We found that Sall1a activates the transcription of key genes involved in hair cell differentiation, and represses genes associated with early sensory fate. Together, these results provide new insights into the molecular basis of hearing impairment in TBS and identify potential therapeutic targets for HC regeneration strategies.

## 2. Results

### 2.1. Sall1a is expressed in inner ear HC after pou4f3 and gfi1.1

To characterize the role of *sall1a* during inner ear development, we first assessed its spatiotemporal expression pattern by in situ hybridization (ISH). We found that *sall1a* is not detectable within the otic vesicle at 30 hours post fertilization (hpf) (Fig1. A-A’). Instead, *gfi1,* a transcription factor implicated in HC differentiation is already expressed at 30 hpf (Fig1. B-B’). When *sall1* ISH is performed in embryos of the reporter line, Tg(pou4f3:gap43-GFP)^s356t^ (Xiao et al., 2005), that labels membranes of HCs, only GFP fluorescence (*pou4f3*) is detected together with *gfi1* and *bmp4* (Fig.1 A-C’). At 48 hpf, *sall1a* expression is detected in differentiating HCs of the anterior and posterior macula (am and pm) (Fig.1 D-G’). At 72 hpf, expression in HCs of both maculae is still visible and expression in lateral cristae starts to appear, as well as in neuromasts HCs (Fig. 1 H-K’). Sall1a is also detected outside the otic vesicle, presumably in otic neurons of the statoacoustic ganglion (SAG). To confirm it, we performed ISH in the reporter line Tg(isl2b:GFP) that labels in green otic neurons. At 30 hpf and 72 hpf, *sall1a* is detected only in a few neurons within the SAG, observed with a larger GFP staining (Fig.1 L-N’). The other paralog, *sall1b* is not expressed in differentiated hair cells but in the sensory progenitors (Suppl. Fig. 1 B-H’). While *sall3* is not detected in the inner ear at the stages analysed (Suppl. Fig. 1 C-J’), *sall4* is expressed in the inner ear but not in HCs (Suppl. Fig. 1 D-L’). Altogether, our data reveals that *sall1a* is expressed after the activation of the differentiating transcription factors, Gfi1.1 and Pou4f3, suggesting that it might act at later stages.

**Figure 1.**
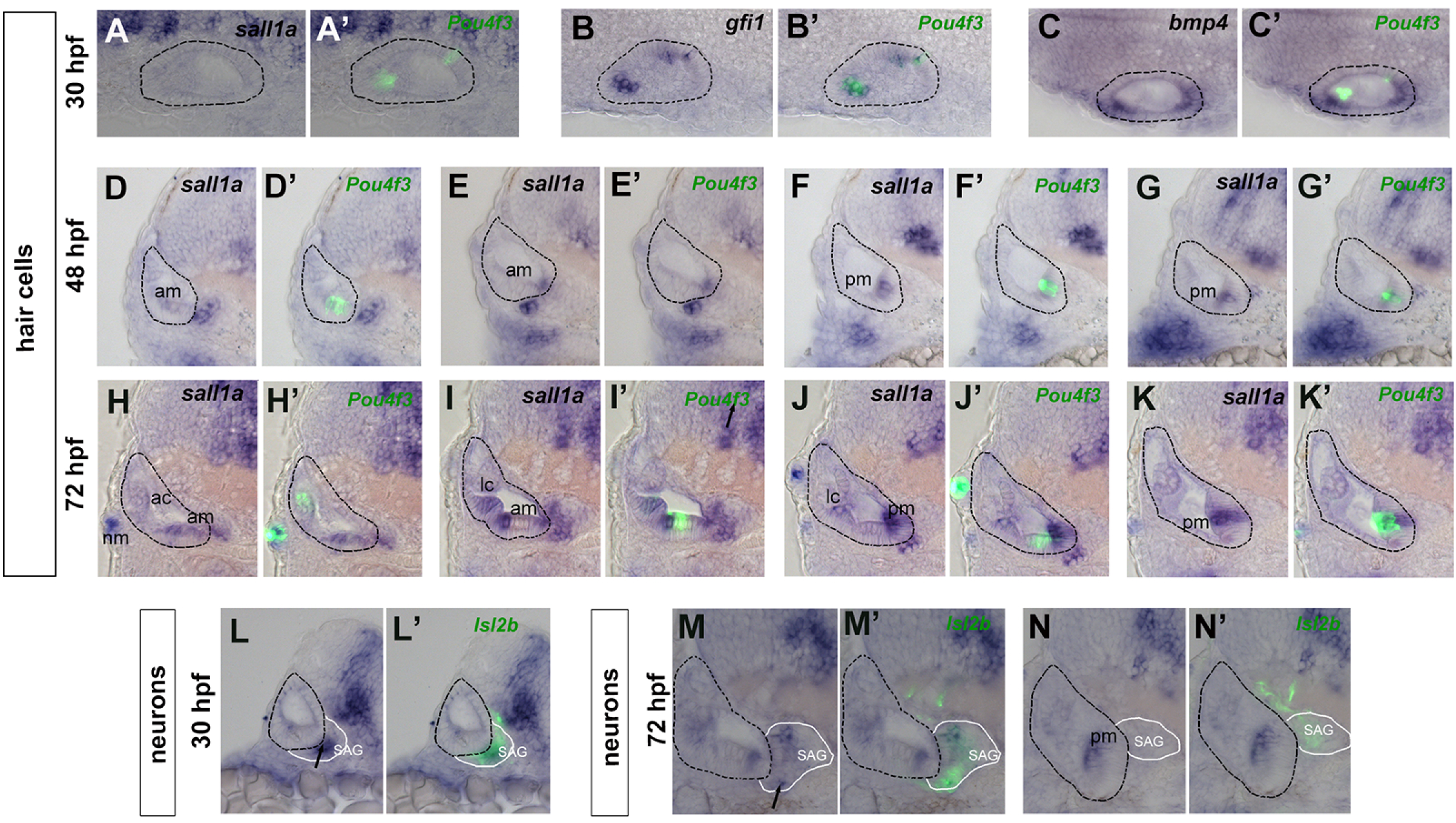
Spatiotemporal expression of *sall1a* mRNA in the otic vesicle of zebrafish embryos. (A–N’) In situ hybridization (ISH) on transverse cryosections of the otic vesicle in transgenic *Tg(pou4f3:gap43-GFP)* zebrafish, where GFP (green) labels hair cells. (A–C’) Expression patterns at 30 hpf showing *sall1a* (A, A’), *gfi1* (B, B’), and *bmp4* (C, C’). (D–G’) Expression of *sall1a* at 48 hpf in hair cell regions (D, F) co-localized with GFP signals. (H–K’) Expression of *sall1a* at 72 hpf showing localization across sensory patches and neruomast. (L-L’) Expression of *sall1a* at 30 hpf colocalizing with islet2b in SAG (black arrows M-N’). No expression is observed at 72 hpf in the SAG. The otic vesicles are outlined with a black dashed line. AC: anterior crista, LC: lateral crista, AM: anterior macula, PM: posterior macula, NM: neuromast, SAG: statoacoustic ganglion.

### 2.2. Sa*ll1a* mutant embryos have a significant reduced number of metabolically active hair cells

To assess the role of *sall1a* in HC development, we have generated a Cas9 KO mutant by injecting at 1-cell stage two CRISPR/Cas9 gRNA homologous to sequences in exon 2 of *sall1a* gene, flanking the region encoding most of its Zinc Finger DNA-binding domains in the HC reporter line described before (Fig. 2A-B, red arrows). These guides also flank the mutation hotspot homologous to the region found in the human ortholog SALL1, responsible for causing TBS. We selected and reared the offspring of two founders and from the first founder, we selected a mutation, consisting in a 2595 bp deletion that ablates 6 out of 9 ZNF domains in the translated protein without causing a downstream frameshift, which results in a protein of 448 amino acids, instead of the 1313 amino acids of the canonical Sall1a protein (Fig. 2C). *Sall1a* KO was confirmed by in situ hybridization assay, at 60 hpf, showing robust expression in WT and heterozygous embryos, but absence in homozygous embryos (Suppl. Fig. 2). We then analyzed the survival rates of descendants from WT crosses and *sall1^+/-^* crosses since larvae homozygous mutants for *sall1a* do not reach adulthood. The survival of WT embryos from 1 and 6 day post-fertilization (dpf) was high (85-90%), while in the offspring of *sall1a^+/-^* mates, the survival dropped to 45-50% (Fig. 2D). We found that homozygous and heterozygous mutants display a lack of stationary balance and larvae cannot maintain a dorsal up position at 5 days post-fertilization (dpf) (Video 1). This phenotype was used to select putative mutant larvae before genotyping.

**Figure 2.**
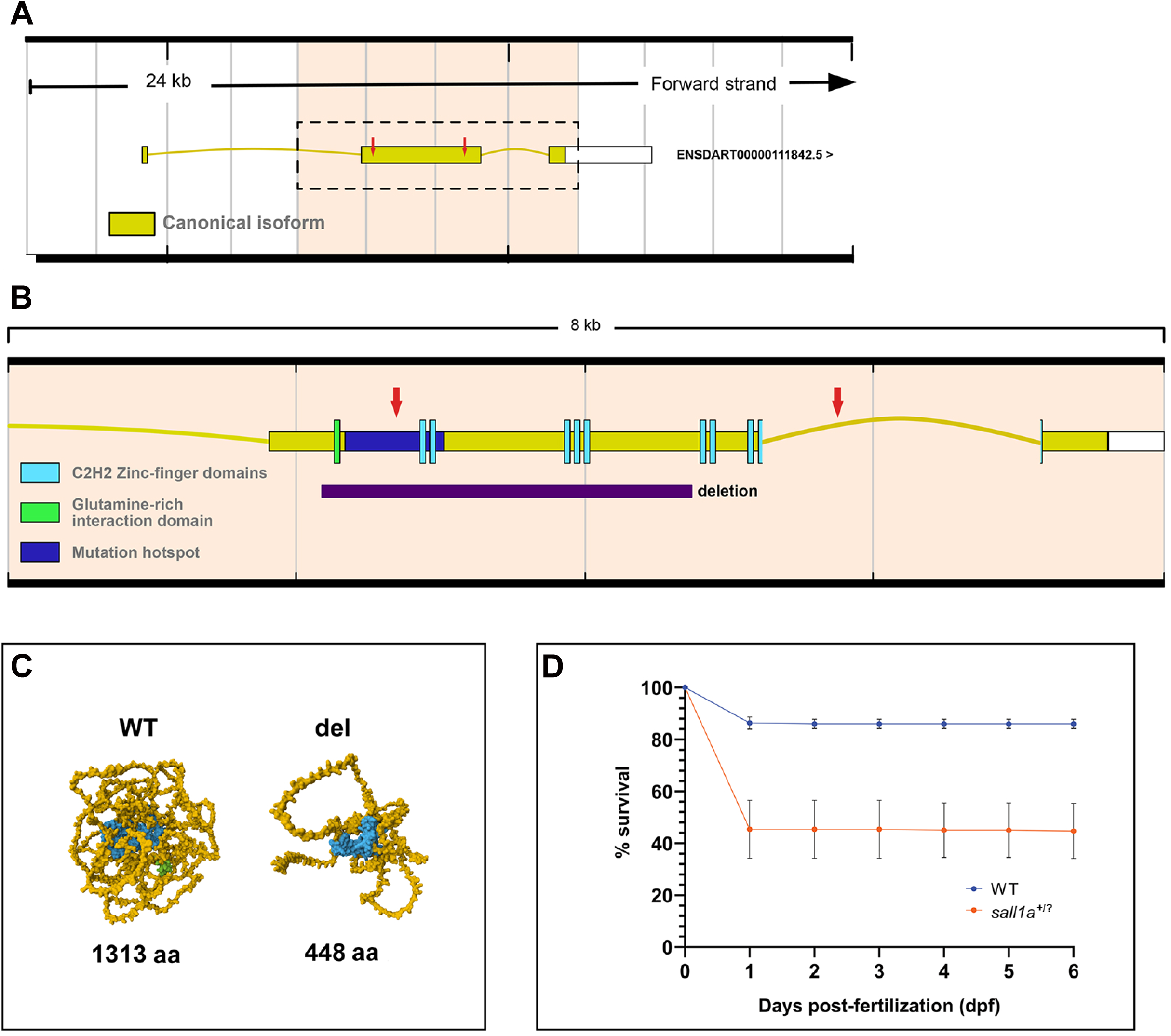
Selected mutation and survival of *sall1a* larvae. (A) Transcript for sall1a gene in zebrafish canonical isoform in yellow (B) Inset of exon 2 from the sall1a canonical isoform and its domains. The glutamine-rich interaction domain is depicted in green, all 9 C2H2 zinc-finger domains are coloured in cyan, and the homologous human mutation hotspot is represented in blue. Selected deletion is shown in purple. (C) Predicted structure of the sall1a WT and mutant proteins. The WT proteins models have been retrieved from the Alphafold Protein Structure Database and the mutant sall1a models have been generated using the Alphafold Server. aa: amino acids (D) Survival curve of sall1a^+/-^ and *sall1a*^-/-^ (orange) and WT (blue) larvae from 0 to 6 dpf (n wt= 300; n sall1a^+/-^, *sall1a*^-/-^ and WT= 300 from 12 independent crosses). *sall1a* larvae come from crosses between two sall1a^+/-^

To study the viability and maturation of HCs at 4 dpf and 6 dpf larvae from *sall1a^+/-^* parents were treated with 4-Di-2-ASP, a vital mitochondrial marker that labels active HCs, and imaged up to eight cranial neuromasts from the lateral line (Fig. 3A-C’). After imaging of neuromasts, individual larvae were genotyped. We found that cranial neuromasts from *sall1a^+/-^* and *sall1a^-/-^* larvae showed a significantly reduction in 4-Di-2-ASP labelled HCs relative to the total number of HCs compared to controls at both 4 dpf (Fig.3 D-F) and 6 dpf (Fig. 3 G-I). Nevertheless, the total number of HCs was not affected in any case. Notably, while the number of 4-Di-2-ASP+ cells was not significantly different in mutant larvae compared to controls at 4 dpf (Fig. 3 E), it was drastically reduced by 6 dpf (Fig. 3 H).

**Figure 3.**
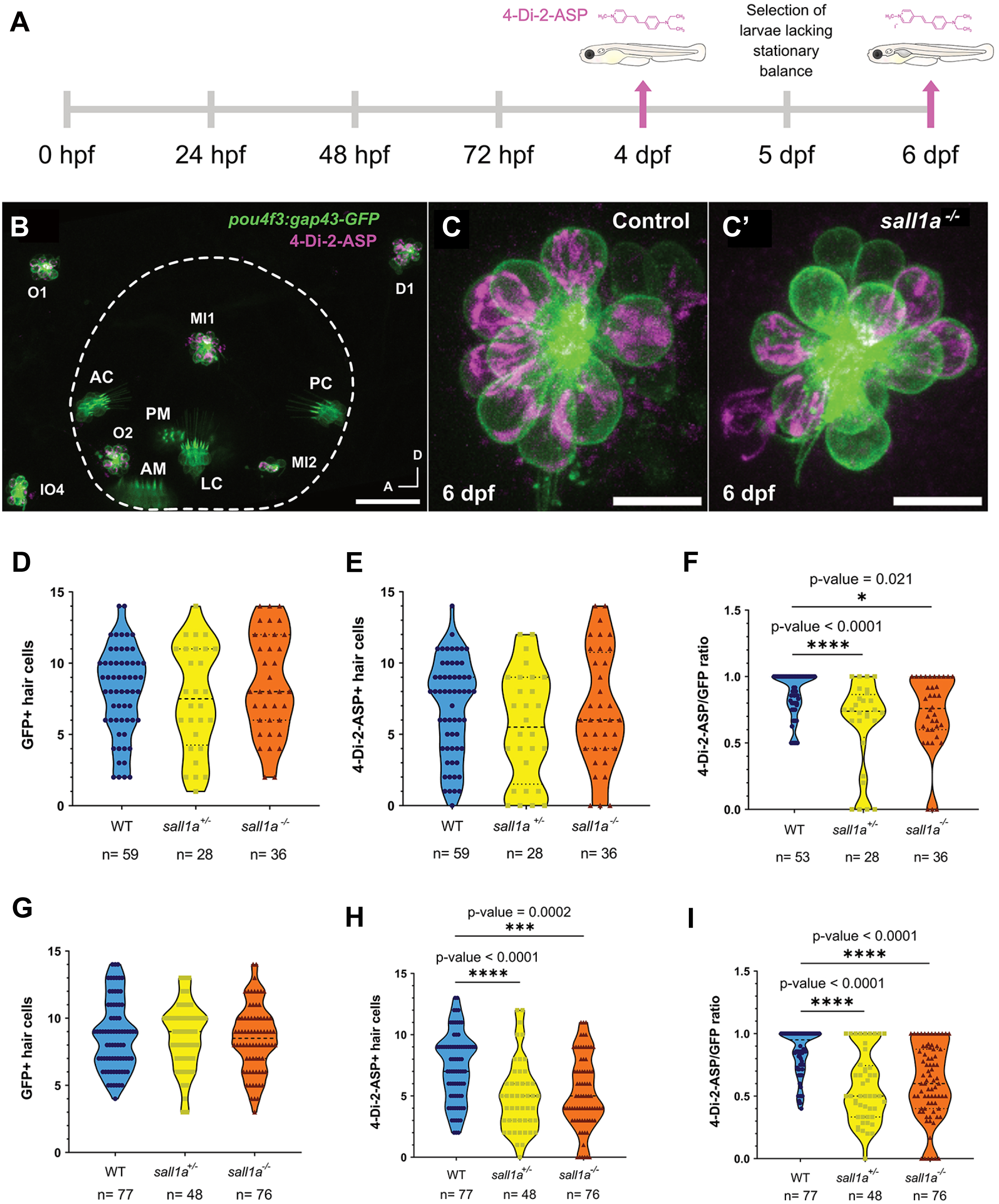
*sall1^-/-^* larvae display a reduced number of metabolically active hair cells. (A) Experimental setup for assessing HC functionality in *sall1a^-/-^* larvae. Control*, sall1a^+/-^*and *sall1a^-/-^* larvae were treated with 4-Di-2-ASP at 4 dpf and 6 dpf and subsequently imaged *in vivo* at the confocal microscope. (B) Confocal maximum Z-projection of the inner ear HC patches (AC, AM, LC, PC and PM) and the surrounding neuromasts from the lateral line (O1, O2, IO4, Ml1, Ml2 and D1) of a larvae treated with 4-Di-2-ASP (magenta) at 6 dpf. Scale bar= 50 µm. Inset of a cranial neuromast from a 6dpf control (C) and a sall1a*^-/-^* embryo (C’) stained with 4-Di-2-ASP. Scale bar = 10 µm. (D-I) Violin plots of GFP+ (D, G), 4-Di-2-ASP+ (E, H) and 4-Di-2-ASP/GFP ratio (F, I) in neuromast HC at 4 dpf (D-F) and 6 dpf (G-I) control and sall1a*^-/-^* larvae. Statistical testing: Kruskal-Wallis test. *p-value≤0.05, **p-value≤0.01, ***p-value≤0.001, ****p-value≤0.0001. AC, anterior crista; AM, anterior macula; PC, posterior crista; LC, lateral crista; PM, posterior macula

These results indicate that the HCs of neuromasts of *sall1a* mutant larvae, whether in heterozygous or homozygous manner, have a reduced number of mature HCs.

### 2.3. sall1a KO larvae exhibit impaired stationary balance, absence of response to tapping stimuli and a rightward turning bias

To investigate whether inhibition of HCs maturation in *sall1a^-/+^*and *sall1a^-^*^/-^ larvae resulted in behavioural defects, we conducted a behavioural testing. Homozygous mutant larvae also exhibited longer periods of inactivity compared to controls but responded to tactile stimuli by initiating movement when the tip of their tail was touched with a pipette tip (Video 2). To further analyse this behaviour and assess vestibular function in the sall1a^-/-^ mutants, 6 dpf larvae were subjected to short-latency (2-12 ms) tapping stimuli using the DanioVision behavioural chamber.

WT and sall1 KO larvae were recorded at 6 dpf during both the absence (habituation) and presence (tapping) of stimuli. Repeated tapping stimulation was applied to a 48-well plate with individualised larvae for a short period (30 seconds) (Fig. 4A), analysing only the first 10 seconds of the presentation, as larvae exhibited a decreasing response to tapping over time. The tapping stimulus activates the zebrafish auditory, vestibular, lateral line, and somatosensory systems, providing a straightforward and effective method for assessing HC function in zebrafish. Larvae were genotyped after the experiment, which confirmed the phenotypic segregation based on the lack of stationary balance in both heterozygous and homozygous mutants. Under physiological conditions, WT zebrafish larvae react to tapping stimuli by exhibiting an acoustic-vibrational startle reflex, characterized by a rapid escape or “C-bend” in response to water-borne mechanical signals.

**Figure 4.**
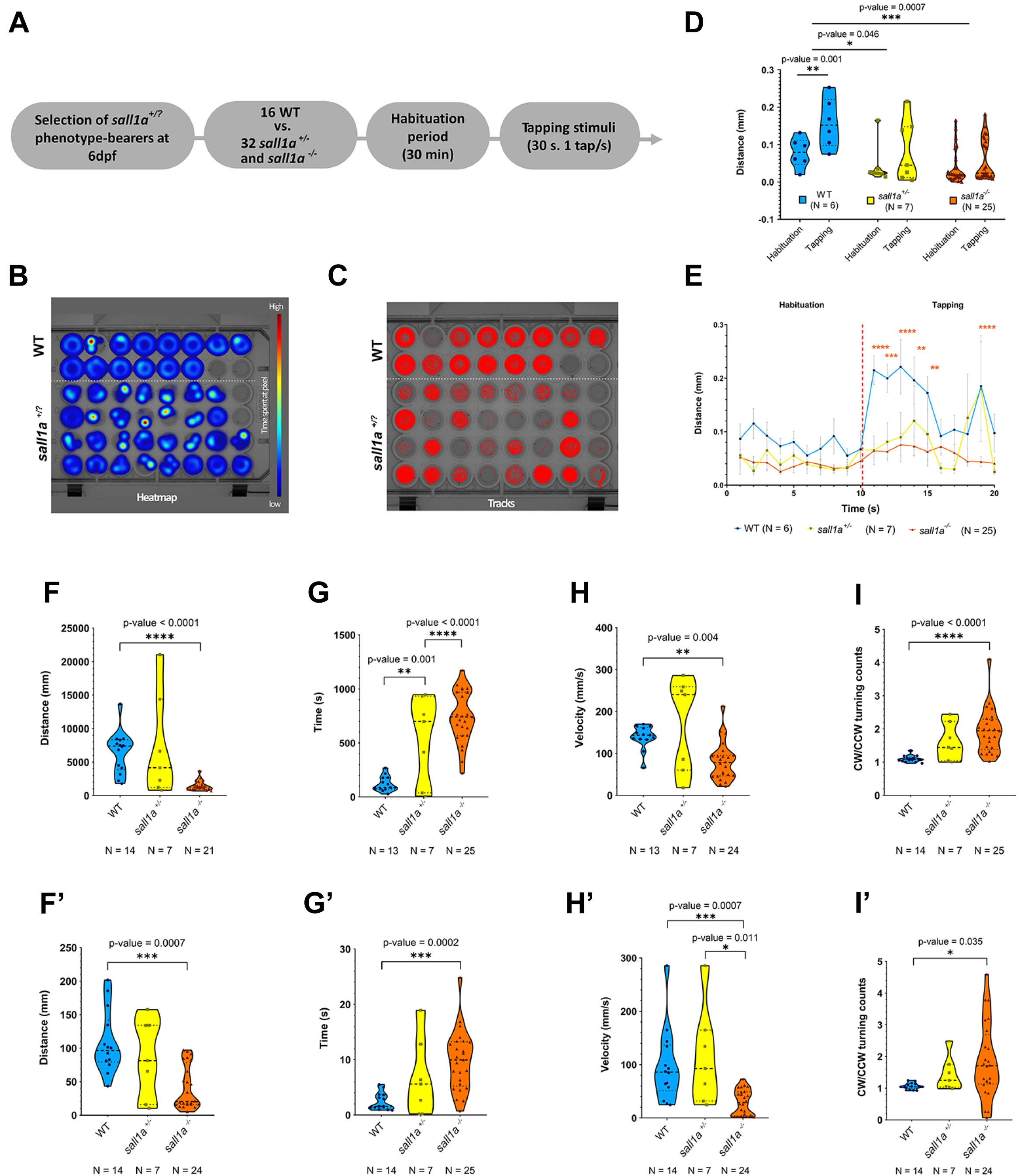
Behavioural analysis of control, *sall1a^+/-^* and *sall1a^-/-^* at 6 dpf. (A) Experimental setup of the behavioural experiments performed with DanioVision (B) Heatmap of the recording. Colder colours in the scale represent a lower amount of time that the larvae have stayed on a pixel, while hotter colours represent a higher amount of time. Embryos with faulty tracking were discarded for further analysis. (C) Tracking of the path (red) followed by larvae on their corresponding wells during the entire experiment. (D) Grouped violin plots comparing the average distance moved per second during the last 10 seconds of habituation compared to the first 10 seconds of tapping. Statistical testing: Two-way ANOVA. (E) Line plot of the average distance moved during the last 10 seconds of habituation compared to the first 10 seconds of tapping. Multiple t-test comparisons of each time point were performed in relation to the WT group. The colour of the significance asterisks (*) matches that of the genotype group with significant changes compared to the WT. (F-H’) Violin plots of the parameters measured during the entire duration of the experiment (F-I) and during tapping (F’-I’): total distance moved (F-F’), total time of inactivity (G-G’), Maximum velocity (H-H’) and ratio of turns (I-I’). Statistical testing: Kruskal-Wallis test. *p-value≤0.05, **p-value≤0.01, ***p-value≤0.001, ****p-value≤0.0001.

Mutant larvae remained inactive for longer periods within their wells and exhibited shorter tracks compared to controls (Fig. 4B and C). ANOVA testing revealed that the observed variation in larval activity was influenced by both the absence/presence of stimuli (p-value= 0.003) and the genotype group (p-value= 0.001) (Fig. 4D). Additionally, Sidak’s post hoc test showed that only WT larvae exhibited a significant increase in the swum distance when comparing the habituation stage to the initial presentation of tapping stimuli. In contrast, mutant larvae with the *sall1a* deletion showed no significant changes between these stages. Both heterozygous and larvae demonstrated a significant decrease in mean distance swum compared to the WT group (Fig. 4D).

A graphical representation of the last 10 seconds of habituation followed by the 10 first seconds of tapping stimuli presentation shows that mutant larvae had an inferior movement profile compared to the WT. Mutants also displayed a very mild reaction to tapping, with significant differences observed between controls and *sall1a^-/-^*larvae at time points when controls demonstrated stronger startle reflex responses (Fig. 4E). Finally, four distinct parameters (total distance swum, total time of inactivity, maximum velocity and ratio of turns) were analysed for both the entire duration of the recording (Fig. 4F-I) and the tapping period specifically (Fig. 4F’-I’). In all cases, *sall1a^-/-^* larvae exhibited significant differences compared to WT. Mutant larvae showed a decrease in total distance swum indicating a rightward turning bias, whereas control larvae did not turn direction. Although comparisons between WT and heterozygous larvae did not yield statistically significant differences, the mean values for each parameter showed noticeable deviations from controls, closely resembling those of the homozygous group. These data suggest that *sall1a* mutant larvae exhibit significant behavioural alterations, likely linked to the observed defects in HC maturation.

### 2.4. Cilia integrity is conserved in hair cells of the cristae and in otic primary cilia

We have shown that *sall1a^+/-^* and *sall1a^-/-^*larvae have compromised mitochondrial function. To further investigate whether the KO affects HC structural integrity, we assessed the stereocilia and kinocilia morphology. Three parameters: kinocilia length, stereocilia bundle length, and stereocilia width were measured in the cristae HCs of WT, *sall1a^+/-^* and *sall1a^-/-^* larvae (Fig. 5). Kinocilia length was quantified across the three cristae and no significance difference was observed between WT and mutants (Fig. A-A’’’). The only notable finding was a significant decrease in stereocilia bundle length in the anterior crista (AC) compared to controls (Fig. 5 B’-B’’’). Moreover, stereocilia bundle width showed a significant difference between heterozygous and homozygous (Fig. 5 C’). Otherwise, both stereocilia and kinocilia appeared structurally normal in mutant larvae in lateral and posterior crista.

**Figure 5.**
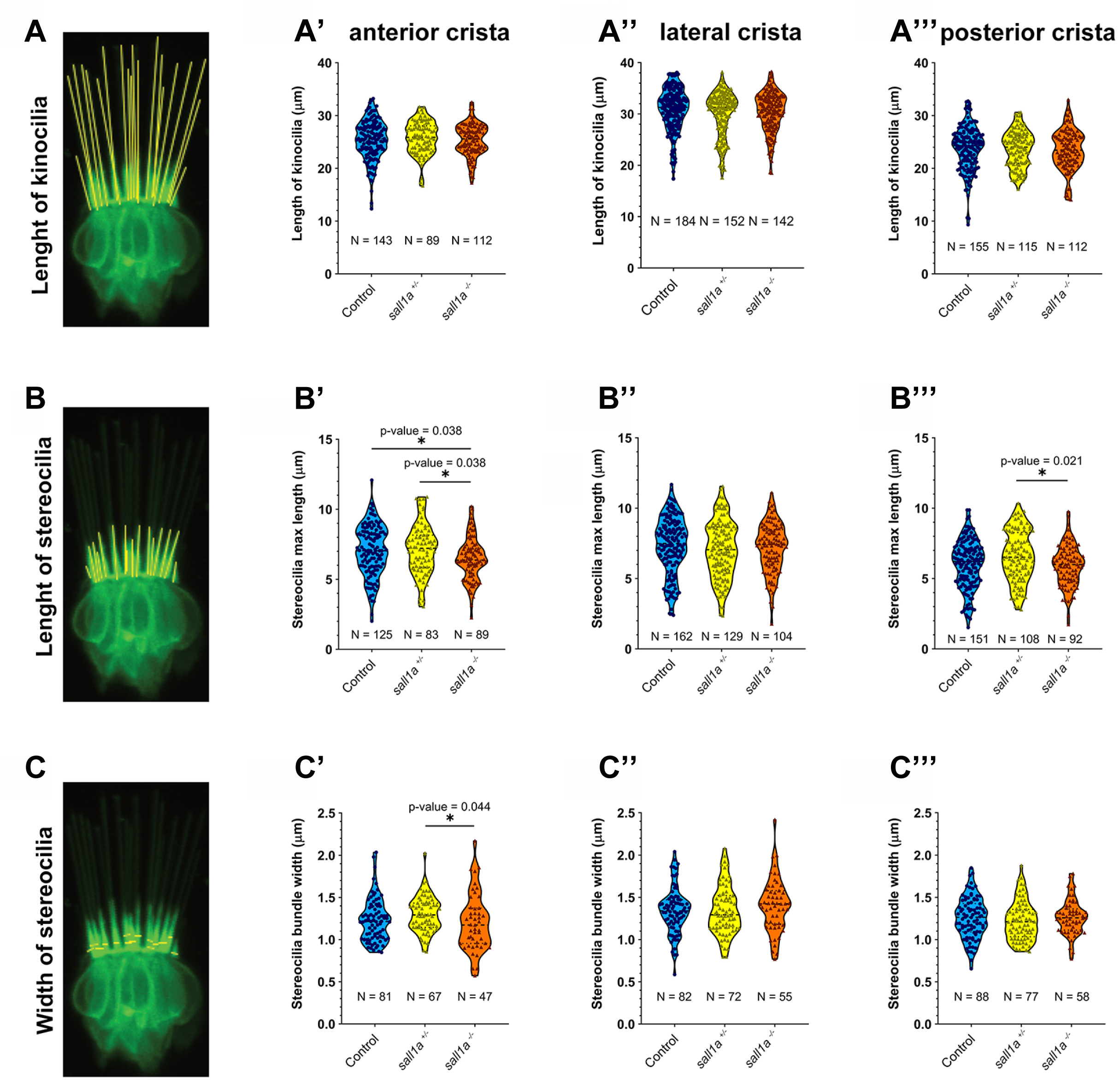
Quantitative analysis of stereocilia and kinocilia of HCs of *sall1a ^+/-^, sall1a ^-/-^* and wild-type larvae. Analysis of kinocilia length in the cristae of the inner ear. (A) Inset of a crista superimposed with kinocilia. Violin plots of the length of kinocilia in HCs from the AC (A’), LC (A’’) and PC (A’’’). (B) Inset of a crista superimposed with stereocilia bundle maximum length. Violin plots of the maximum length of stereocilia bundles in HCs from the AC (B’), LC (B’’) and PC (B’’’). (C) Inset of a crista superimposed with stereocilia bundle width. Violin plots of the width of stereocilia bundles in HCs from the AC (C’), LC (C’’) and PC (C’’’). Statistical testing: Mann-Whitney (2 group comparison) and Kruskal-Wallis tests (> 2 groups). *p-value < 0.05. Scale bar = 20 µm.

### 2.5. RNA-seq of hair cells from *sall1a* mutant larvae reveals downregulation of hearing loss genes and significant differential expression in mitochondrial related genes

To identify direct targets of *sall1a*, we performed a transcriptomic analysis in purified HCs by Fluorescent Activated Cell Sorting from 60 hpf embryos from a cross of heterozygous adults (only 12 hours after its induction) and WT embryos. We identified 271 differentially expressed (DE) genes (Fig. 6C-D). Each of these genes was individually examined, and information regarding their fold change expression was documented (Suppl. Table 1). These genes were then categorized based on their relevance to inner ear and HCs developmental pathways. From this group, 15 genes were selected for further analysis, including three downregulated genes associated with SNHL in humans and expressed in mice and zebrafish HCs (Blasiole et al., 2006; Kazmierczak et al., 2017) (Fig. 6F). Following the differential expression analysis, we conducted a functional enrichment analysis of the DE genes, examining Gene Ontology (GO) and Kyoto Encyclopedia of Genes and Genomes (KEGG) pathways. For GO pathways, we generated a ranked list of biological processes (BP) and identified eight over-represented BP GO terms with an adjusted p-value < 0.05. All eight BP terms were related to mitochondria, including terms like ATP synthesis coupled electron transport and oxidative phosphorylation. These findings align with the observed phenotype of reduced HC maturity (Fig. 6), which is closely associated with mitochondrial volume and architecture due to the high energy demands of HCs. The KEGG pathway analysis also highlighted oxidative phosphorylation as an over-represented pathway, with most of the associated genes being upregulated. When extending the analysis to include GO terms with an adjusted p-value beyond 0.05, additional terms emerged, such as nervous system development (adjusted p-value = 0.057) and posterior lateral line (pLL) development (adjusted p-value = 0.125), which include genes like *pou4f2*, *fgf3, pax6a*, *isl2b*, *isl1*, *irx4a*, *sox2* or *atoh1a*. Specifically, downregulation of *fgf3* in the anterior macula of the inner ear in homozygous *sall1a* mutant embryos was confirmed by in situ hybridization (Suppl. Fig. 3).

**Figure 6.**
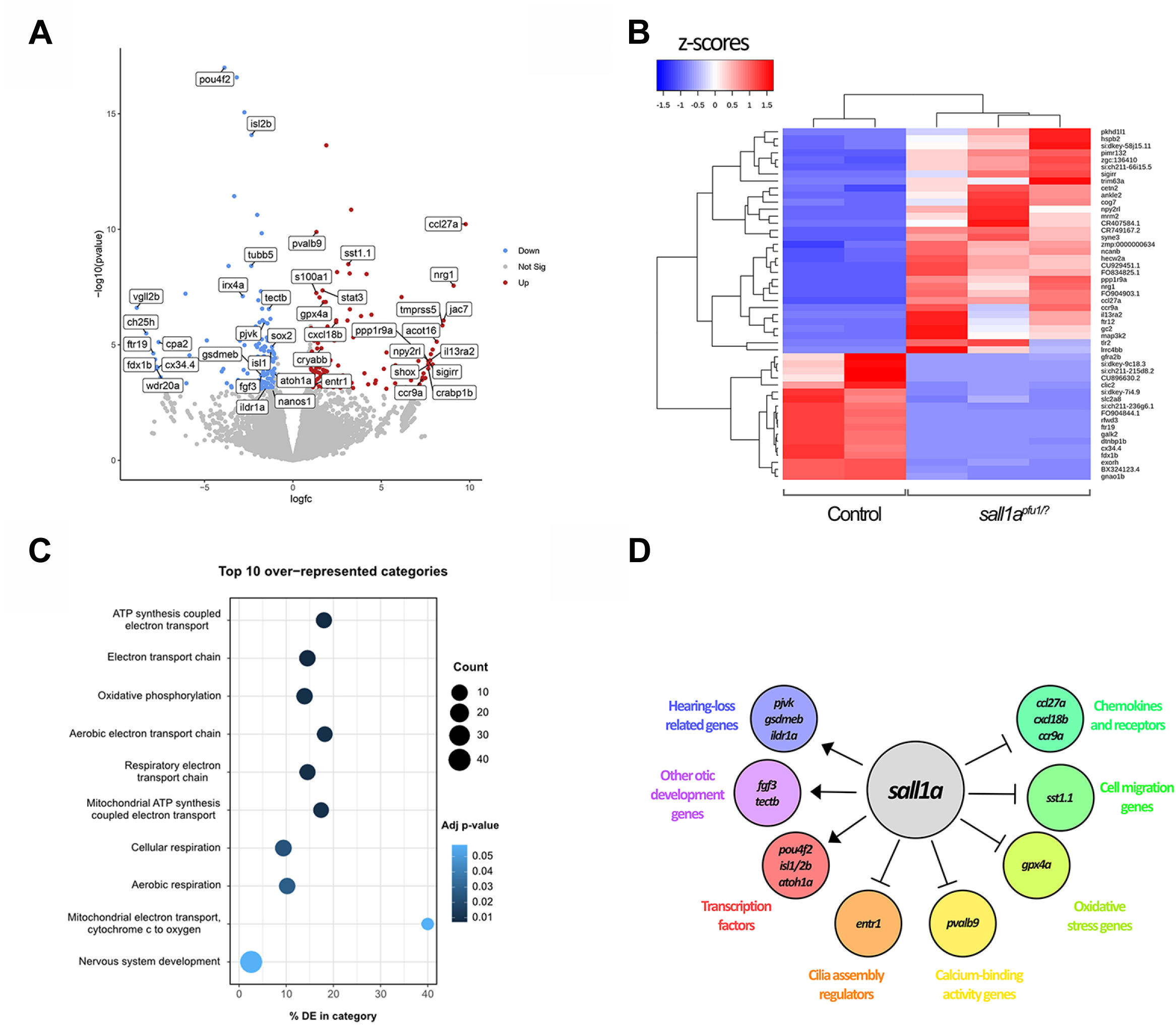
RNA-seq of *sall1a* HCs reveals differentially expressed genes related to hearing loss disorders and a disruption of mitochondria-related pathways. (A) Volcano plot of differentially expressed (DE) genes. Grey: non-significant, Blue: Downregulated in the *sall1a^+/?^* group, Red: Upregulated in the *sall1a^/?^* Group. 40 DE genes related to HC development were labelled. (B) Heatmap of Z-scores for the top 50 DE genes with a standard deviation lower than 25. Blue: Downregulated genes, Red: Upregulated genes. (C) Top 10 over-represented categories for GO terms of biological pathways. (D) Schematic representation of the interaction of the transcription factor sall1a with some selected DE genes, grouped by function.

## 3. Discussion

HC maturation is closely linked to acquiring functionality, which includes developing structures for sensing mechanical stimuli, opening mechanotransduction channels to depolarize cells, maintaining a robust mitochondrial network to meet high energy demands, and effectively communicating stimuli to sensory neurons in the form of electrical signals (Caprara and Peng, 2022; Gillespie and Müller, 2009). Mutations in the human *SALL1* gene on chromosome 16q12.1 cause Townes-Brocks syndrome (TBS), a syndrome associated with developmental abnormalities, including sensorineural hearing loss (Wang and Streit, 2024). However, the molecular mechanisms underlying the inner ear phenotype remain poorly understood. Here, we aimed to elucidate the role of Sall1a in HC development to provide insights into the molecular mechanisms underlying hearing loss in TBS.

To fulfil this objective, we generated a *sall1a* KO transgenic zebrafish line and performed molecular and behavioral analysis. Our results identified a previously unrecognized role for Sall1a as a transcriptional regulator that acts as a maturation factor of HCs. Importantly, *sall1a* expression begins after the onset of previously identified differentiation transcription factors such as *Pou4f3* and *Gfi1.1* (Hertzano et al., 2004; Keithley et al., 1999). This temporal pattern indicates that Sall1a is not involved in early fate specification (before 48 hpf) but during the transition from differentiated to mature HCs (after 48 hpf). Furthermore, *sall1a* mutants exhibited markedly reduced survival compared with WT, highlighting the impact of loss of *sall1a* and suggesting that its functions may extend beyond HCs maturation. In humans, heterozygous deletions of the entire SALL1 gene have been associated with milder manifestations of TBS, while homozygous SALL1 mutations are incompatible with embryonic viability (Borozdin et al., 2006).

Consistent with this, the total number of HCs in neuromasts was not significantly altered in *sall1a* mutant larvae. However, the proportion of 4-Di-2-ASP positive HCs was reduced in both heterozygous and homozygous larvae, suggesting a defect in HC maturation or mitochondrial activity. As 4-Di-2-ASP enters HCs through open MET channels of stereocilia (Zhu et al., 2021), the differences observed during maturation reflect changes in the proportion of HCs with open MET channels. The reduction of 4-Di-2ASP positive cells between 4 and 6 dpf reinforces the idea that Sall1a is required for maturation rather than for their initial differentiation. The cellular phenotype was accompanied by clear behavioral abnormalities. Interestingly, *sall1a* mutant larvae showed impaired stationary balance and reduced startle response at 6 dpf, which correlates with vestibular affectations. Intriguingly, the *sall1a* mutant embryos also displayed a rightward turning bias. These results suggest that *sall1a* deficiency affects sensory functions required for postural control. Although the phenotypes observed in zebrafish larvae cannot be directly compared to TBS signs, they support the potential role of SALL1 in sensory system function.

To further assess whether behavioural defects might be due to a phenotype in the ciliary structures, we examined their morphology. The maximum length of the stereocilia bundle and kinocilium can serve as proxies for HCs age, as both structures elongate during maturation (McQuate et al., 2023). At 6 dpf, kinocilia length and stereocilia width were preserved, however a modest reduction in stereocilia bundle length in the AC was observed, thus the phenotype cannot be attributed to large morphological defects. Altogether, these results suggest that the integrity of ciliary structures is preserved in *sall1a* mutant larvae. However, this does not rule out the possibility that mechanotransduction may still be impaired, which could explain the described stationary balance phenotype. Further studies may help to elucidate whether the mechanosensory machinery is affected at later stages. However, at 60 hpf, none of the major components of the mechanotransduction channels were found to be directly downregulated in our transcriptional analysis. Our results contrast with those of *in vitro* models of TBS, where marked elongation of primary cilia has been reported, highlighting that truncated SALL1 proteins alter ciliary transport dynamics (Bozal-Basterra et al., 2018). This can be explained by the fact that only the truncated Sall1 protein acting as a dominant-negative is responsible for interacting with ciliary proteins.

In addition to investigating the morphological, functional, and behavioural changes in *sall1a* mutant larvae to understand the role of this transcription factor in inner ear development, we also explored its potential transcriptional activation or repression of primary target genes in HCs. To this end, we analysed gene expression in HCs from both control and mutant larvae at 60 hpf, before the onset of the previously mentioned phenotypic alteration. RNA-seq revealed 271 differentially expressed genes between both conditions. Among them, *pjvk*, *gsdmeb, and ildr1a* that are associated with sensorineural hearing loss (Borck et al., 2011; Domínguez-Ruiz et al., 2022) were downregulated. Our DE gene list also included relevant HC genes such as *isl2*b, *atoh1a* and *fgf3*. This suggests that under physiological conditions, *sall1a* may function as a transcriptional activator for these genes. In addition, over 30 mitochondrial-associated genes cause human hearing loss, as HCs produce approximately 75% of their ATP via oxidative phosphorylation (McQuate et al., 2023). Our KEGG pathway analysis also highlighted oxidative phosphorylation as an over-represented pathway, but surprisingly most of the associated genes were upregulated. Although upregulation of oxidative phosphorylation genes is typically associated with increased ATP production and improved mitochondrial function, suggesting a higher degree of maturation (O’Sullivan et al., 2023), this might not always be the case. Maturation issues in HCs that compromise mitochondrial function may trigger the activation of the mitochondrial transcriptional machinery to address the dysfunction. Altogether, our findings position Sall1 as a key maturation factor of HCs and provide novel insights of its function in inner ear development, which might elucidate its role in hearing loss associated with SALL1 mutations in TBS humans.

## 4. Materials and Methods

### 4.1. Fish maintenance and husbandry

Zebrafish (Danio rerio) embryos and adults were maintained and handled according to standard procedures (Westerfield, 2007) at the aquatic facility of the Parc de Recerca Biomèdica de Barcelona, in compliance with the guidelines of the European Community Directive (2010/63/EU) and the Spanish legislation for the experimental use of animals. For this study we used the Tg(pou4f3:gap43-GFP)s356t (Xiao et al., 2005) to visualize hair cells from the inner ear and the lateral line. Embryos used later than 24 hpf were kept transparent by soaking them in 1X Danieau’s solution with 1X N-phenylthiourea (PTU) (Sigma-Aldrich, P7629) to inhibit tyrosinase activity and subsequent pigment formation (Westerfield, 2007). Embryos were incubated at 28.5 °C in the dark, staged as described (Kimmel et al., 1995) and fixed at the desired stage in 4% paraformaldehyde (PFA) (Sigma-Aldrich, P6148) in 1X phosphate buffered saline (PBS) pH 7.5 for 3 hours at RT or 4°C overnight (O/N). For in vivo experiments requiring anaesthesia, larvae were immersed in 1X Danieau’s solution containing MS-222 (Sigma-Aldrich, A5040) at a final concentration of 200 mg/L.

### 4.2. CRISPR guide RNA design and microinjection

Novel transgenic mutant line was generated with CRISPR Cas9-mediated genome editing to generate a stable knockout line (Tg(pou4f3:gap43-GFP)s356t;sall1apfu1/+). crRNAs (supplied at 2 nM) and tracrRNA (supplied at 20 nM) were obtained from Integrated DNA Technologies (IDT). The efficacy of crRNAs was previously evaluated in silico using CHOPCHOP, CRISPRscan and CRISPOR. The selected crRNA sequences targeting the exon 2 are: CAGTTCCAACACCACTGAGT abd GCTACGGTAATGGATGTCCA. The 36-nucleotide crRNA was resuspended in 20 μL of the IDTE buffer to obtain a final concentration of 100 μM. The 67-nucleotide tracrRNA was resuspended in 200 μL of the IDTE buffer to obtain a final concentration of 100 μM. For RNA duplex formation, crRNA and tracrRNA (1:1) were combined with IDT Duplex Buffer and subjected to thermal annealing. 1.5 µL resulting RNA duplex (C_f_ = 5 µM) was combined with 1.5 µL of recombinant Ca9 protein (C_f_ = 3 µM) to form the RNP complex. 1-2 nL of the assembled RNP complexes were injected into the cell in one-cell-stage embryos using an Eppendorf Femtojet 4i microinjector.

### 4.3. Genotyping of sall1a deletion

To identify large deletions of embryos from the *sall1a* transgenic lines, fin clipping and PCR genotyping was realized with the design of three primers: one forward (Fw1) and two reverse primers (Rv1 and Rv2). The forward primer (Fw1) was designed to anneal upstream of the deletion site, while the reverse primers were positioned within the deleted region (Rv1) and downstream of the deletion (Rv2) (Figure SX, supplementary material). PCR protocol was as follows: Fw1 ATACTTGACAAAGAGGAGGCCA primer, Rv1 TGAGGTAGTGAGGCAGAGATGA primer and Rv2 ATCATAGAGGACAAGGGTCCAA 35 cycles of 94°C 3 min; 96°C 10 s; 57°C 15 s; 72°C 30 s; 72°C 10 min; hold at 4°C. Following amplification, 5 μL of each PCR product, mixed with 10X FastDigest Green Buffer (ThermoFisher Scientific, B72), was loaded into a 3% agarose gel for electrophoresis. three distinct banding patterns were observed. In WT embryos, only a 179 bp band produced by the Fw1-Rv1. Conversely, in homozygous mutant embryos, only a 312 bp band amplified by the Fw1-Rv2 primer pair was present, as the deletion prevented Rv1 from annealing to the gDNA. Finally, heterozygous embryos displayed both bands: a 179 bp band from the WT allele and a 312 bp band.

### 4.4. 4-Di-2-ASP *in vivo* staining

This protocol was based on the guidelines provided by (Nakae et al., 2012). Briefly, a 10 mM 4-Di-2-ASP stock solution was prepared in dimethylformamide and stored at 4C. For staining, the stock solution was diluted in 1x Danieau’s solution, final concentration of0.1 mM. Zebrafish embryos were incubated in the staining solution for at least 30 min with gentle agitation in the dark, followed by multiple washes in 1x Danieau’s solution to remove excess dye. Prior to imaging, embryos were anesthetized with MS-2222 (200 mg/L) and prepared for confocal microscopy.

### 4.5. Confocal imaging

For visualization of the zebrafish inner ear, embryos were typically mounted laterally, ensuring that one otic vesicle was in close contact with the glass bottom of a 35 mm Ø glass-bottom culture dish (MatTek, P35G-1.5-14-C). Embryos were carefully dechorionated using forceps and placed in 1% low-melting agarose (Ecogen, AG0400) dissolved in distilled water, initially heated to 70°C and then maintained at 37°C to prevent premature solidification. The agarose solution contained 200 mg/L of tricaine to anesthetize the embryos and minimize movement during in vivo imaging. Imaging was conducted using a Leica TCS SP8 confocal microscope (Leica Microsystems). Frames were captured at a resolution of 1024×512 with an 8-bit depth. The primary objective used for most experiments was an HC PL APO 20x/0.70 IMM CORR immersion objective (Leica Microsystems, #506191 – discontinued product) set in the glycerol position. The scan speed was maintained at 600 Hz, and Z-step sizes were optimized using Leica system settings to avoid oversampling. The HC PL APO 20x/0.70 IMM CORR immersion objective typically had an optimized Z-step size of 1.19 μm. A software zoom of 1.5 to 2.5x was applied in most experiments, with bidirectional scanning enabled to reduce acquisition time. Line averaging was set to 1 or 2 to enhance image quality.FIJI software version 2.14.0/1.54f was used for image processing.

### 4.6. In situ hybridization

Riboprobes were generated by *in vitro* transcription using DIG dNTPs of cloned cDNA or directly from PCR-amplified fragments, using primers containing the promoter sequence of T7 or SP6 sequences at the 5’ ends of the reverse primer.

Embryos were fixed at the desired stage and dehydrated with methanol and stored at - 20°C until the use. After rehydration, permeabilization with proteinase K (diluted 1:1000 from a stock of 10 mg/mL, Invitrogen) during 5-30 mins according to the stage. Embryos were washed for 3×5min with PBST 0.1% and postfixed for 30 mins with PFA 4%. After fixation, larvae were washed 2 x 5 mins with PBST 0.1% and incubated with the hybridization buffer (50% FAD, 25% SSC, 250 mg torula RNA, 1.25 mg heparin, Tween 0.1%) at 70°C for 2h. Incubate ON at 70°C with the riboprobes diluted in hybridization buffer at a concentration of 0.5 ng/µL (DIG-labeled probes). After 2×30 min washes with 50%FAD/2xSCC, 1×15 min wash with 2xSCC and 1×30 min wash 0.2 SCC buffer, 4×15 min washes with MABT (malic acid buffer in PBT) an incubation with blocking solution (2% blocking reagent (Roche), 10% neutralized goat serum in 1x MABT) was done for 1h. Incubation with anti-DIG-AP (1:2000), was performed O/N at 4°C. After several washes in MABT, color reaction with NBT/BCIP was performed. The staining time ranged from 1h to 2 days, probe-dependent. After stopping the reaction with dH_2_O, embryos were kept in PBST/Glycerol 1:1 until imaging.

### 4.7. Behavioural test and analysis

Behavioural tests were conducted using the DanioVision behavioural chamber (Noldus Information Technology) under constant illumination (100% intensity, ∼4400 lux) with a capture rate of 25 frames per second. Individual larvae were placed in the wells of a 48-well plate, each containing 1 mL of embryo with water. A total of 16 control larvae were tested alongside 32 mutant larvae at 6 dpf, selected based on their inability to self-right. Prior to testing, the plate was incubated for 1 hour at 28.5°C under light conditions to allow for habituation after transport to the experimental room.

Upon placement in the DanioVision chamber, the larvae underwent an additional 30-minute acclimatization at 28.5°C under light conditions. The experiment consisted of two phases: an initial 30-minute free-swimming period followed by a startle response test. The startle response was elicited by 30 mechanical taps. applied at 1-second intervals during the final 30 seconds of the trial. These taps were generated using a solenoid positioned at the base of the DanioVision chamber, with the stimulus intensity set to the highest level (level 8). Behavioural tracking was performed using Noldus EthoVision XT software (Noldus Information Technology), employing dynamic subtraction as the detection method. Larvae with poor tracking data were excluded from the final analysis. The following behavioural parameters were analysed during both the free-swimming and startle response phases: average distance moved, total distance moved, maximum distance moved, total inactivity time, average velocity, maximum velocity, and turning angles. Following the behavioural assay, the *sall1a* mutant larvae were genotyped to confirm the presence of the mutation, distinguishing between heterozygous and homozygous genotypes.

### 4.8. Sample preparation for RNA sequencing

GFP-positive embryos were selected at 48 hpf. Three biological replicates were prepared for each condition (sall1a ^+/-^ *, sall1a^-/-^* and control groups). Embryos were dechorionated, anesthetized, and dissected at 60 hpf to remove tissue anterior to the otic vesicles. The embryos were then transferred to a 4-well plate for dissociation in preparation for cell sorting. After removing the water, 600 μL of pre-warmed protease solution (Trypsin 1X, EDTA 1mM, PBS 1X) at 28.5°C was added to the embryos, followed by pipetting for 15 minutes. Following dissociation, stop solution (Fetal Bovine Serum 30%, CaCl2 6mM and PBS 1X) was added and samples were centrifuged at 200 g for 10 minutes. Cells were resuspended in 500 μL of Medium 199 (Merck, M4530-100ML) supplemented with fetal bovine serum. DAPI was added to cell suspension to assess the viability and fluorescence-activated cell sorting (FACS) of GFP-positive cells was performed. FACS was performed using a BD Influx cell sorter (BD Biosciences, 23-10536-00) using a 0.35 μm filter in single-drop mode. Up to 1000 GFP-positive cells were collected per sample in 0.2 mL tubes containing freshly prepared CDS Sorting Solution (10X Lysis buffer, Rnae inhibitor, 3’ SMART-Seq CDS Primer IIA, Water). RNA extraction and Cdna synthesis from the sorted cells were performed by using the SMART-Seq® v4 Ultra® Low Input RNA Kit for Sequencing (Takara, 634888). The samples were briefly spun and incubated at 72°C for 3 minutes in a hot-lid thermal cycler. A Reverse Transcription Master Mix was prepared, and after a 2-minute ice incubation, 7.5 μL of the Reverse Transcription Master Mix was added to each sample. cDNA synthesis was conducted at 42°C for 90 minutes, followed by 70°C for 10 minutes and a 4°C hold O/N. cDNA was amplified, PCR Master Mix was prepared, and 30 μL were added to each tube containing 20 μL of first-strand cDNA. The thermal cycler program included an initial denaturation at 95°C for 1 minute, followed by 13 cycles of 98°C for 10 seconds, 65°C for 30 seconds, and 68°C for 3 minutes, with a final extension at 72°C for 10 minutes. Amplified cDNA was stored at 4°C O/N. Amplified cDNA was purified using magnetic bead-based purification. AMPure XP beads (Beckman Coulter, A63880).The quality and quantity of the cDNA were assessed using a 4200 Tapestation (Agilent, G2991BA) with a High Sensitivity D5000 ScreenTape® assay (Agilent, 5067-5592, 5067-5593 & 5067-5594). Libraries were prepared using the Nextera XT DNA Library Prep Kit (Illumina, 15032354 & 15032355). DNA tagmentation was performed by mixing 5 μL of Tagment DNA Buffer with 2.5 μL of input DNA (1 ng) and 2.5 μL of Amplicon Tagment Mix, followed by incubation at 55°C for 7 minutes. The reaction was neutralized with 2.5 μL of Neutralise Tagment Buffer, centrifuged at 280g at 20°C for 1 minute, and incubated at RT for 5 minutes.

Library amplification was achieved by adding 2.5 μL each of i7 and i5 adapters (Illumina, 15028211), along with 7.5 μL of Nextera PCR Master Mix. The samples were subjected to the following thermal cycler program: 72°C for 3 minutes, denaturation at 95°C for 30 seconds, 13 cycles of 95°C for 10 seconds, 55°C for 30 seconds, and 72°C for 30 seconds, followed by a final extension at 72°C for 5 minutes. AMPure XP beads were used for library cleanup. Following centrifugation and 25 μL of supernatant transfer to new tubes, 15 μL of beads were added, and the mixture was incubated for 5 minutes at RT. After bead binding, the tubes were placed on a magnetic separation device for 2 minutes and the resulting supernatant was discarded. The beads were washed twice with 200 μL of fresh 80% ethanol while on the magnetic separation device. Residual ethanol was removed, and the beads were air-dried for 15 minutes before eluting the DNA with 26.25 μL of RSB buffer. The tubes were briefly centrifuged again, followed by a 2-minute incubation at RT. The tubes were then placed back on the magnetic separation device until the liquid became completely clear. A total of 50 μl of supernatant, containing the eluted DNA, was carefully transferred to new 1.5 mL tubes. the library quality was verified using a 4200 Tapestation with a High Sensitivity D1000 ScreenTape® assay (Agilent, 5067-5584, 5067-5585, 5067-5587 & 5067-5603) to confirm size distribution (250-1,000 bp). The final libraries were sequenced using the NextSeq 500 platform (Illumina, SY-415-1001) with a 2×75 cycles mid-output (Illumina, FC-404-1001) run, yielding approximately 120 million reads in total, with 20 million reads per sample.

### 4.9. RNA-sequencing analysis

RNA-seq analysis was conducted on the European server of the Galaxy platform, as described previously (Batut et al., 2021). The analysis pipeline included several key steps: data quality control, mapping to the zebrafish reference genome, read quantification per annotated gene, differential gene expression analysis, and functional enrichment of DE genes. Paired-end data were analysed using the FastQC tool, and results were aggregated with MultiQC for streamlined visualization. After the initial quality assessment, data were filtered and trimmed with the Cutadapt tool, removing paired reads if either was shorter than 38 bp or longer than 2000 bp, with a quality cut-off of 30. Next, reads were mapped to the zebrafish reference genome (GRCz11) using RNA STAR. Post-mapping checks were conducted to ensure data integrity, including assessing read duplication levels (MarkDuplicates), mapping statistics per chromosome (Samtools idxstats), gene body coverage (by subsampling 200,000 reads with Samtools view), and read distribution across features (Read Distribution tool on the RNA STAR output). Read quantification began with an estimation of strand specificity to determine whether mapped reads originated from forward or reverse-encoded transcripts, which is crucial for reads overlapping genes on opposite strands. Our data were determined to be unstranded, as confirmed through STAR coverage visualization in pyGenometracks and read subsampling using the Infer Experiment tool. Reads were then counted for each annotated gene using featureCounts. Differential gene expression analysis was performed using DESeq2, which normalizes for sequencing depth and library composition. ‘Condition’ was set as the primary factor, with two levels: ‘Knockout’ and ‘Control,’ and all comparisons were made relative to the Control group. Quality control included generating a PCA plot, a heatmap of sample-to-sample distances, and an MA-plot. DESeq2 generated a list of differentially expressed (DE) genes, which were annotated with Ensembl identifiers using the "Annotate DESeq2/DEXSeq output tables" tool. Genes were then filtered based on a p-value < 0.05 and a fold change greater than 2 or less than 0.5, yielding 271 DE genes. Z-scores of normalized counts were calculated and visualized in a heatmap using the heatmap2 tool, and DE genes were additionally visualized in a volcano plot. Finally, functional enrichment analysis of the DE genes was performed using GO and KEGG pathway analysis via the goseq and Pathview tools.

### 4.10. Statistical analysis

All data were initially tested for normal distribution using the Kolmogorov-Smirnov test, and homogeneity of variances was assessed with Levene’s test. For comparisons between two groups, a two-tailed Student’s t-test was applied to parametric data when populations exhibited equal standard deviations. In cases where equal standard deviations could not be assumed, Student’s t-test with Welch’s correction was employed. For non-parametric data, the Mann-Whitney U test was utilized. When comparing more than two groups, one-way ANOVA followed by an appropriate post hoc test was used for parametric data. For non-parametric comparisons, the Kruskal-Wallis test was conducted. Graphs were generated using GraphPad Prism 8.0.1 software, with error bars representing the SEM.

## Authorship contribution statement

**Nerea Montedeoca**: Writing – original draft, Investigation, Formal analysis, Methodology, Data curation. **Francesca Manocchio**: Writing – original draft, review & editing, Investigation, Methodology. **Laura Fargas**: Formal analysis, Methodology. **Berta Alsina**: Conceptualization, Writing – review & editing, Data Curation, Project administration, Funding acquisition.

## Declaration of competing interest

The authors declare that they have no competing interests.

## Funding

This work was supported by Ministerio de Ciencia e Innovación PID2023-146798NB-I00, PID2020-117662GB-I00 (FEDER) to Berta Alsina. Nerea Montedeoca was recipient of the predoctoral fellowship from funds of Excelencia María de Maeztu Award, funded by the AEI (CEX2018-000792-M).

## Acknowledgment

We thank members of the Neurobiology of Sensory Systems group at UPF and the CRG-ALMU microscopy facility staff for technical support in image acquisition in SP8 microscopy.

## Abbreviations

4-Di-2-ASP: 4-(4-Diethylaminostyryl)-1-methylpyridinium iodide;
Aa: amino acids;
AC: Anterior Crista;
Dpf: days post-fertilization;
EGFP: Enhanced Green Fluorescent Protein;
FACS: Fluorescence-activated cell sorting;
FISH: Fluorescent In Situ Hybridization;
GO: Gene Ontology;
hpf: hours post-fertilization;
PC: Posterior Crista;
pLL: Posterior Lateral Line;
PM: Posterior Macula;
SALL: Spalt-like;
TBS: Townes-Brocks syndrome,
WT: wild-type.

## Notes

### Competing Interest Statement

The authors have declared no competing interest.

### Summary of Updates

Reference section: improved Authors: Laura Fargas Madriles added

