## Supplementary Figures for "Sall1a, a transcription factor responsible for the Townes-Brocks syndrome, is involved in hair cell maturation"

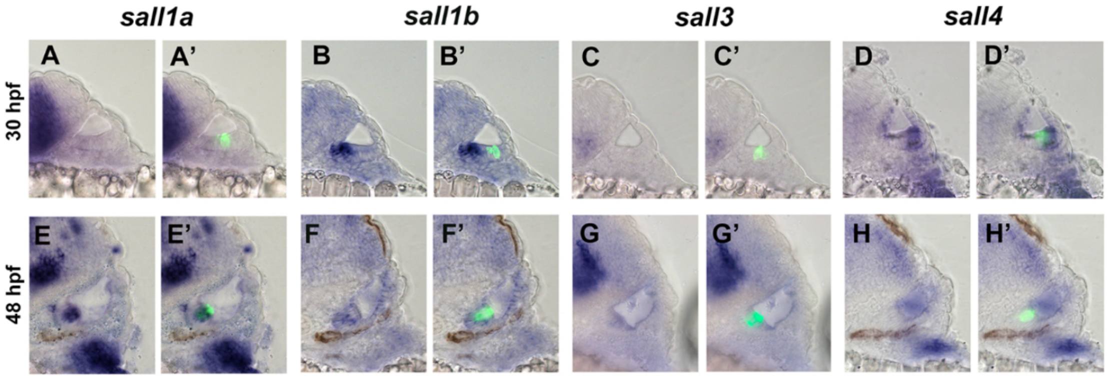


**Supplementary Figure 1. Spatiotemporal expression of the mRNA of the spalt like family members in the otic vesicle of zebrafish embryos.**
In situ hybridization of *sall1a*, *sall1b, sall3* and *sall4* on transverse cryosections of the inner ear in zebrafish larvae with a Tg(pou4f3: gap43 GFP background at 30 hpf (A-D') and 48 hpf (E-H').

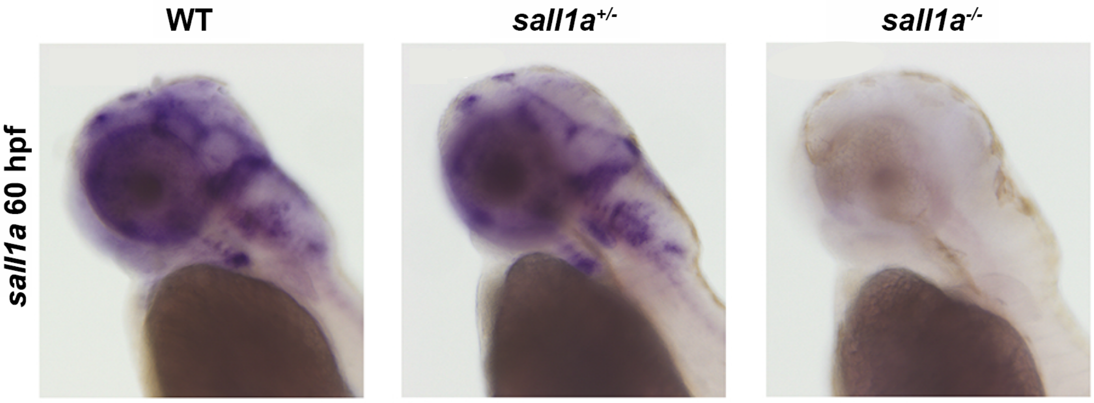


**Supplementary Figure 2. Expression of sall1a at 60 hpf.**
Validation of knockout with in situ hybridization of sall1a at 60 hpf in wt*, sall1a*$^{+/-}$ *and sall1a*$^{-/-}$*.*

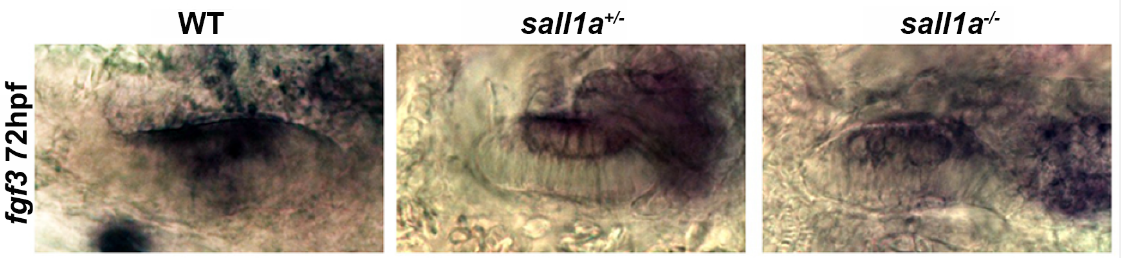


**Supplementary Figure 3. Expression of *fgf3* in zebrafish embryos.**
Validation of RNA-seq data with in situ hybridization of *fgf3* of wt, sall1a$^{+/-}$ and sall1a$^{+/-}$ embryos in the otic vesicle at 72 hpf.
