## Supplementary material for "Sall1a, a transcription factor responsible for the Townes-Brocks syndrome, is involved in hair cell maturation": Suppl Table

| P-value | P-adj | log2(FC) | Fold change | Abs(FC) | Status |
|---|---|---|---|---|---|
| 1,53496E-41 | 1,42076E-37 | 5,518097028 | 45,82608164 | 45,82608164 | Upregulated |
| 2,37089E-08 | 1,09725E-05 | 2,650428887 | 6,278539003 | 6,278539003 | Upregulated |
| 6,04348E-09 | 3,02706E-06 | 2,613531722 | 6,120000294 | 6,120000294 | Upregulated |
| 8,89749E-12 | 1,02944E-08 | 2,607375678 | 6,093941616 | 6,093941616 | Upregulated |
| 3,13432E-09 | 1,74961E-06 | 2,40662452 | 5,302322865 | 5,302322865 | Upregulated |
| 3,21341E-09 | 1,74961E-06 | 2,381570628 | 5,21103747 | 5,21103747 | Upregulated |
| 5,50077E-07 | 0,000145472 | 2,300326628 | 4,92569271 | 4,92569271 | Upregulated |
| 1,26804E-06 | 0,000255151 | 2,288070167 | 4,884023584 | 4,884023584 | Upregulated |
| 1,15634E-07 | 3,82255E-05 | 2,241112195 | 4,727613828 | 4,727613828 | Upregulated |
| 2,46967E-07 | 7,14353E-05 | 2,233429149 | 4,702503929 | 4,702503929 | Upregulated |
| 5,31537E-08 | 2,04996E-05 | 2,159170719 | 4,466580363 | 4,466580363 | Upregulated |
| 1,06311E-06 | 0,000227459 | 2,125864772 | 4,364646377 | 4,364646377 | Upregulated |
| 4,31878E-06 | 0,000726811 | 2,100216714 | 4,287737882 | 4,287737882 | Upregulated |
| 6,21371E-09 | 3,02706E-06 | 2,073371462 | 4,208690622 | 4,208690622 | Upregulated |
| 1,62635E-06 | 0,000313615 | 1,960525176 | 3,892036326 | 3,892036326 | Upregulated |
| 4,94758E-07 | 0,000134691 | 1,936805345 | 3,82856923 | 3,82856923 | Upregulated |
| 1,00049E-06 | 0,000220488 | 1,911156688 | 3,761105274 | 3,761105274 | Upregulated |
| 2,44085E-06 | 0,000461072 | 1,894462678 | 3,717834829 | 3,717834829 | Upregulated |
| 4,49519E-06 | 0,000729955 | 1,813706128 | 3,515442088 | 3,515442088 | Upregulated |
| 3,43479E-14 | 5,29874E-11 | 1,769154542 | 3,408541486 | 3,408541486 | Upregulated |
| 7,04867E-05 | 0,006154955 | 1,7479141 | 3,358725983 | 3,358725983 | Upregulated |
| 0,000208425 | 0,01349076 | 1,713712767 | 3,28003854 | 3,28003854 | Upregulated |
| 9,83987E-07 | 0,000220488 | 1,664009758 | 3,168960677 | 3,168960677 | Upregulated |
| 3,90997E-05 | 0,003951512 | 1,639159841 | 3,11484385 | 3,11484385 | Upregulated |
| 1,95293E-07 | 5,86443E-05 | 1,627334453 | 3,089416655 | 3,089416655 | Upregulated |
| 0,000654733 | 0,028318723 | 1,599352874 | 3,030073677 | 3,030073677 | Upregulated |
| 0,000674076 | 0,028885425 | 1,579961202 | 2,989618097 | 2,989618097 | Upregulated |
| 0,000394352 | 0,020166425 | 1,568862588 | 2,966707286 | 2,966707286 | Upregulated |
| 1,21412E-07 | 3,87513E-05 | 1,558056722 | 2,944569493 | 2,944569493 | Upregulated |
| 0,000227223 | 0,014380655 | 1,544250086 | 2,916524283 | 2,916524283 | Upregulated |
| 0,000599039 | 0,026879868 | 1,530085485 | 2,888029512 | 2,888029512 | Upregulated |
| 3,6978E-08 | 1,61012E-05 | 1,523191676 | 2,874262202 | 2,874262202 | Upregulated |
| 0,000235244 | 0,014419967 | 1,516106265 | 2,860180639 | 2,860180639 | Upregulated |
| 0,000114538 | 0,008908931 | 1,509024984 | 2,846176212 | 2,846176212 | Upregulated |
| 0,000475843 | 0,02269432 | 1,47810741 | 2,785830361 | 2,785830361 | Upregulated |
| 0,000244078 | 0,014575418 | 1,473893635 | 2,777705472 | 2,777705472 | Upregulated |
| 0,000103446 | 0,008399084 | 1,470626566 | 2,771422309 | 2,771422309 | Upregulated |
| 0,001389304 | 0,047918152 | 1,462770587 | 2,75637196 | 2,75637196 | Upregulated |
| 0,001035831 | 0,040513847 | 1,45724061 | 2,745826767 | 2,745826767 | Upregulated |
| 0,00048702 | 0,022882545 | 1,44815865 | 2,728595712 | 2,728595712 | Upregulated |
| 0,000480563 | 0,02269432 | 1,446385687 | 2,725244534 | 2,725244534 | Upregulated |
| 0,000127347 | 0,009589296 | 1,419438076 | 2,674813077 | 2,674813077 | Upregulated |
| 0,001150155 | 0,042926753 | 1,413022967 | 2,662945618 | 2,662945618 | Upregulated |
| 9,85266E-05 | 0,008076631 | 1,407616457 | 2,652984887 | 2,652984887 | Upregulated |
| 1,00001E-05 | 0,001358548 | 1,406342918 | 2,650643998 | 2,650643998 | Upregulated |
| 9,20075E-08 | 3,27547E-05 | 1,3845141 | 2,610840094 | 2,610840094 | Upregulated |
| 0,000810219 | 0,033479419 | 1,381119622 | 2,60470434 | 2,60470434 | Upregulated |
| 0,000407213 | 0,020373871 | 1,349938682 | 2,549012913 | 2,549012913 | Upregulated |
| 0,000181058 | 0,012182963 | 1,339572056 | 2,530762381 | 2,530762381 | Upregulated |

| | | | | | |
|---|---|---|---|---|---|
| 0,000232845 | 0,014380655 | 1,297668204 | 2,458312298 | 2,458312298 | Upregulated |
| 7,00944E-05 | 0,006154955 | 1,297571849 | 2,458148117 | 2,458148117 | Upregulated |
| 1,08127E-06 | 0,000227459 | 1,289795455 | 2,444933889 | 2,444933889 | Upregulated |
| 6,70196E-10 | 4,77179E-07 | 1,286664765 | 2,439634064 | 2,439634064 | Upregulated |
| 0,000139665 | 0,009888229 | 1,284533192 | 2,436032182 | 2,436032182 | Upregulated |
| 3,1123E-05 | 0,003200825 | 1,276672733 | 2,422795661 | 2,422795661 | Upregulated |
| 7,73918E-06 | 0,001102059 | 1,274925522 | 2,419863252 | 2,419863252 | Upregulated |
| 0,000535733 | 0,024903573 | 1,269685407 | 2,411089838 | 2,411089838 | Upregulated |
| 4,03827E-10 | 3,39802E-07 | 1,267483351 | 2,407412481 | 2,407412481 | Upregulated |
| 0,001164684 | 0,043121267 | 1,25761049 | 2,390993968 | 2,390993968 | Upregulated |
| 1,03811E-07 | 3,5588E-05 | 1,240915892 | 2,3634853 | 2,3634853 | Upregulated |
| 1,32128E-05 | 0,001675304 | 1,240357878 | 2,362571314 | 2,362571314 | Upregulated |
| 0,000762638 | 0,031797205 | 1,235110539 | 2,35399383 | 2,35399383 | Upregulated |
| 0,000547617 | 0,025092772 | 1,23442989 | 2,352883501 | 2,352883501 | Upregulated |
| 0,00068611 | 0,029124848 | 1,222692186 | 2,333818206 | 2,333818206 | Upregulated |
| 0,000141016 | 0,009888229 | 1,220534246 | 2,33032996 | 2,33032996 | Upregulated |
| 0,000226549 | 0,014380655 | 1,21554414 | 2,322283557 | 2,322283557 | Upregulated |
| 0,001122258 | 0,042525682 | 1,213487806 | 2,318975866 | 2,318975866 | Upregulated |
| 0,001499434 | 0,049621495 | 1,208584048 | 2,311106987 | 2,311106987 | Upregulated |
| 0,001483196 | 0,049613154 | 1,205299296 | 2,305851007 | 2,305851007 | Upregulated |
| 0,000233049 | 0,014380655 | 1,191725519 | 2,28425786 | 2,28425786 | Upregulated |
| 8,68386E-05 | 0,007307077 | 1,175419486 | 2,258585433 | 2,258585433 | Upregulated |
| 7,25332E-07 | 0,000181451 | 1,159530126 | 2,233846612 | 2,233846612 | Upregulated |
| 4,04199E-05 | 0,003980071 | 1,145200786 | 2,211769107 | 2,211769107 | Upregulated |
| 0,000448761 | 0,021747293 | 1,131760562 | 2,191259828 | 2,191259828 | Upregulated |
| 0,00086563 | 0,035297761 | 1,122155603 | 2,176719642 | 2,176719642 | Upregulated |
| 6,80034E-06 | 0,00099911 | 1,118950473 | 2,171889152 | 2,171889152 | Upregulated |
| 0,000294186 | 0,016705441 | 1,11580772 | 2,167163079 | 2,167163079 | Upregulated |
| 0,000133528 | 0,009808734 | 1,108127658 | 2,155657027 | 2,155657027 | Upregulated |
| 3,0682E-05 | 0,003190923 | 1,101496978 | 2,145772279 | 2,145772279 | Upregulated |
| 0,001283451 | 0,046045043 | 1,091069971 | 2,130319726 | 2,130319726 | Upregulated |
| 0,000427285 | 0,021036958 | 1,090949367 | 2,130141646 | 2,130141646 | Upregulated |
| 0,000315831 | 0,016996124 | 1,080012857 | 2,114054921 | 2,114054921 | Upregulated |
| 0,000186583 | 0,012424537 | 1,076476873 | 2,108879807 | 2,108879807 | Upregulated |
| 0,000188137 | 0,012432485 | 1,061398266 | 2,086953225 | 2,086953225 | Upregulated |
| 0,001240257 | 0,045288261 | 1,058280577 | 2,082448153 | 2,082448153 | Upregulated |
| 6,03641E-05 | 0,005531979 | 1,05618556 | 2,07942631 | 2,07942631 | Upregulated |
| 4,65036E-06 | 0,000742134 | 1,055161926 | 2,07795142 | 2,07795142 | Upregulated |
| 4,74386E-05 | 0,004480532 | 1,051042744 | 2,072026916 | 2,072026916 | Upregulated |
| 0,000239262 | 0,014487125 | 1,049708178 | 2,070111072 | 2,070111072 | Upregulated |
| 0,000424704 | 0,021021691 | 1,04777875 | 2,067344403 | 2,067344403 | Upregulated |
| 0,00030954 | 0,016875017 | 1,047434362 | 2,066850963 | 2,066850963 | Upregulated |
| 0,001089492 | 0,042018068 | 1,045563453 | 2,064172375 | 2,064172375 | Upregulated |
| 0,001137907 | 0,042641553 | 1,030722361 | 2,043046955 | 2,043046955 | Upregulated |
| 0,000342941 | 0,017972809 | 1,021341968 | 2,029806169 | 2,029806169 | Upregulated |
| 0,000127689 | 0,009589296 | 1,016471597 | 2,022965332 | 2,022965332 | Upregulated |
| 0,000617913 | 0,027235255 | 1,014986213 | 2,02088358 | 2,02088358 | Upregulated |
| 0,000245667 | 0,01457623 | 1,006806227 | 2,009457726 | 2,009457726 | Upregulated |
| 2,89583E-05 | 0,003080895 | 1,00351109 | 2,004873333 | 2,004873333 | Upregulated |
| 6,25131E-05 | 0,005672754 | 0,999147823 | 1,998818981 | 1,998818981 | Upregulated |

| | | | | | |
|---|---|---|---|---|---|
| 9,13731E-05 | 0,007619368 | 0,98591426 | 1,980568034 | 1,980568034 | Upregulated |
| 3,34774E-06 | 0,000584654 | 0,974833276 | 1,965414052 | 1,965414052 | Upregulated |
| 0,001281803 | 0,046045043 | 0,966089065 | 1,953537656 | 1,953537656 | Upregulated |
| 0,000148433 | 0,010330022 | 0,962436749 | 1,94859835 | 1,94859835 | Upregulated |
| 0,00147774 | 0,049613154 | 0,961616107 | 1,947490251 | 1,947490251 | Upregulated |
| 0,000602843 | 0,026879868 | 0,958431057 | 1,943195504 | 1,943195504 | Upregulated |
| 0,001119133 | 0,042525682 | 0,958019128 | 1,942640747 | 1,942640747 | Upregulated |
| 0,000117443 | 0,009058745 | 0,953272468 | 1,936259702 | 1,936259702 | Upregulated |
| 2,92869E-06 | 0,000531528 | 0,95169135 | 1,934138825 | 1,934138825 | Upregulated |
| 0,000728592 | 0,030653847 | 0,94287581 | 1,922356369 | 1,922356369 | Upregulated |
| 1,94085E-05 | 0,002273984 | 0,936868013 | 1,914367769 | 1,914367769 | Upregulated |
| 0,000109551 | 0,008723307 | 0,928445775 | 1,903224535 | 1,903224535 | Upregulated |
| 1,56391E-05 | 0,001930074 | 0,926066995 | 1,900088999 | 1,900088999 | Upregulated |
| 1,85536E-05 | 0,002201689 | 0,918720536 | 1,890438002 | 1,890438002 | Upregulated |
| 0,000310387 | 0,016875017 | 0,916677484 | 1,88776278 | 1,88776278 | Upregulated |
| 0,000972526 | 0,038633903 | 0,915270391 | 1,8859225 | 1,8859225 | Upregulated |
| 0,001392608 | 0,047918152 | 0,9055634 | 1,873275903 | 1,873275903 | Upregulated |
| 0,001323182 | 0,047105292 | 0,903970681 | 1,871208967 | 1,871208967 | Upregulated |
| 0,001506474 | 0,049622513 | 0,902664872 | 1,869516069 | 1,869516069 | Upregulated |
| 0,00113022 | 0,042525682 | 0,892615757 | 1,856539174 | 1,856539174 | Upregulated |
| 0,000399549 | 0,020208897 | 0,88124309 | 1,841961734 | 1,841961734 | Upregulated |
| 0,001383057 | 0,047918152 | 0,880359041 | 1,840833369 | 1,840833369 | Upregulated |
| 0,000406279 | 0,020373871 | 0,879108937 | 1,839238967 | 1,839238967 | Upregulated |
| 0,000635261 | 0,027867202 | 0,869164356 | 1,82660458 | 1,82660458 | Upregulated |
| 2,81762E-05 | 0,003032545 | 0,857123808 | 1,81142341 | 1,81142341 | Upregulated |
| 0,001100634 | 0,042271647 | 0,853332435 | 1,806669277 | 1,806669277 | Upregulated |
| 8,43974E-05 | 0,007166806 | 0,846909559 | 1,798643866 | 1,798643866 | Upregulated |
| 0,000469956 | 0,022538423 | 0,844481118 | 1,795618814 | 1,795618814 | Upregulated |
| 0,001501082 | 0,049621495 | 0,833056304 | 1,781455325 | 1,781455325 | Upregulated |
| 0,000411013 | 0,020453441 | 0,813913927 | 1,757974235 | 1,757974235 | Upregulated |
| 0,000311758 | 0,016875017 | 0,804547841 | 1,7465983 | 1,7465983 | Upregulated |
| 0,001419493 | 0,04830451 | 0,785640664 | 1,723857671 | 1,723857671 | Upregulated |
| 0,000308672 | 0,016875017 | 0,779243069 | 1,716230192 | 1,716230192 | Upregulated |
| 0,000663652 | 0,028570969 | 0,775053068 | 1,711252996 | 1,711252996 | Upregulated |
| 0,001088852 | 0,042018068 | 0,750369725 | 1,682223886 | 1,682223886 | Upregulated |
| 0,001350113 | 0,047515748 | 0,716108308 | 1,642744724 | 1,642744724 | Upregulated |
| 0,00148475 | 0,049613154 | 0,692034854 | 1,615560586 | 1,615560586 | Upregulated |
| 3,36774E-18 | 1,55859E-14 | -3,196249461 | 0,109102082 | 9,165727921 | Downregulated |
| 6,5843E-18 | 2,03148E-14 | -2,774716237 | 0,146125895 | 6,843414046 | Downregulated |
| 8,8569E-13 | 1,17113E-09 | -2,696274608 | 0,154290955 | 6,481261339 | Downregulated |
| 7,86137E-10 | 5,19749E-07 | -2,63865575 | 0,160577788 | 6,227511366 | Downregulated |
| 6,97035E-16 | 1,61294E-12 | -2,442825453 | 0,183923094 | 5,437055118 | Downregulated |
| 3,82699E-08 | 1,61012E-05 | -2,208960278 | 0,216290128 | 4,623419528 | Downregulated |
| 1,05663E-14 | 1,95603E-11 | -2,132516688 | 0,22805968 | 4,384817167 | Downregulated |
| 3,31506E-06 | 0,000584654 | -2,072437095 | 0,237757524 | 4,20596573 | Downregulated |
| 2,51376E-09 | 1,55116E-06 | -2,017741882 | 0,246944393 | 4,049494653 | Downregulated |
| 5,04139E-06 | 0,0007909 | -1,975504196 | 0,254281041 | 3,932656546 | Downregulated |
| 5,65898E-10 | 4,36496E-07 | -1,905952702 | 0,266840083 | 3,747562923 | Downregulated |
| 1,40489E-11 | 1,44485E-08 | -1,843424146 | 0,278659618 | 3,588607519 | Downregulated |
| 7,17298E-05 | 0,006204964 | -1,821212264 | 0,282983087 | 3,533780094 | Downregulated |

| | | | | | |
|---|---|---|---|---|---|
| 0,000189389 | 0,012432485 | -1,776068757 | 0,291977933 | 3,424916361 | Downregulated |
| 9,46824E-07 | 0,000219095 | -1,756464203 | 0,295972655 | 3,378690506 | Downregulated |
| 0,000112514 | 0,008825696 | -1,741869567 | 0,298981979 | 3,344683197 | Downregulated |
| 6,31918E-08 | 2,33961E-05 | -1,716854699 | 0,304211227 | 3,287189664 | Downregulated |
| 2,8874E-06 | 0,000531528 | -1,701283515 | 0,307512399 | 3,2519014 | Downregulated |
| 5,57193E-06 | 0,000859563 | -1,701133301 | 0,307544419 | 3,251562829 | Downregulated |
| 0,000268309 | 0,0156259 | -1,691124554 | 0,309685437 | 3,229083066 | Downregulated |
| 7,58762E-06 | 0,001097359 | -1,657586305 | 0,316969009 | 3,154882566 | Downregulated |
| 7,74349E-07 | 0,000188615 | -1,63206555 | 0,322625963 | 3,099564553 | Downregulated |
| 2,12797E-10 | 1,96965E-07 | -1,631001481 | 0,322864006 | 3,097279291 | Downregulated |
| 5,2062E-08 | 2,04996E-05 | -1,618679589 | 0,32563336 | 3,070938432 | Downregulated |
| 0,0002767 | 0,016007097 | -1,612490388 | 0,327033337 | 3,057792243 | Downregulated |
| 6,38533E-06 | 0,000964115 | -1,608705929 | 0,327892333 | 3,049781592 | Downregulated |
| 2,17636E-05 | 0,002518049 | -1,607968547 | 0,328059966 | 3,048223203 | Downregulated |
| 1,9641E-07 | 5,86443E-05 | -1,553878171 | 0,340593269 | 2,936053325 | Downregulated |
| 0,000279606 | 0,016074715 | -1,545285124 | 0,34262798 | 2,918617446 | Downregulated |
| 0,000480355 | 0,02269432 | -1,531667765 | 0,345877299 | 2,891198704 | Downregulated |
| 1,2785E-05 | 0,00164358 | -1,529772643 | 0,346331942 | 2,887403323 | Downregulated |
| 6,11571E-07 | 0,000157242 | -1,524032688 | 0,347712614 | 2,875938227 | Downregulated |
| 4,43197E-06 | 0,000729955 | -1,521104651 | 0,348419035 | 2,870107256 | Downregulated |
| 0,000259968 | 0,015326519 | -1,496683911 | 0,354366981 | 2,821933344 | Downregulated |
| 9,9901E-06 | 0,001358548 | -1,495408985 | 0,354680278 | 2,819440671 | Downregulated |
| 0,001261377 | 0,045753014 | -1,490950831 | 0,355777991 | 2,810741605 | Downregulated |
| 0,00023947 | 0,014487125 | -1,49093422 | 0,355782087 | 2,810709242 | Downregulated |
| 7,03344E-05 | 0,006154955 | -1,48662321 | 0,356846812 | 2,802322917 | Downregulated |
| 7,29298E-05 | 0,006250353 | -1,475610494 | 0,359581202 | 2,781013009 | Downregulated |
| 0,000141002 | 0,009888229 | -1,450903904 | 0,365792169 | 2,733792803 | Downregulated |
| 0,000134584 | 0,009808734 | -1,448480201 | 0,36640721 | 2,729203935 | Downregulated |
| 3,01777E-05 | 0,003174146 | -1,442109068 | 0,368028891 | 2,717177981 | Downregulated |
| 9,32253E-07 | 0,000219095 | -1,440561381 | 0,368423915 | 2,714264623 | Downregulated |
| 0,001416087 | 0,04830451 | -1,419690125 | 0,37379259 | 2,675280427 | Downregulated |
| 1,44492E-05 | 0,001807325 | -1,417291541 | 0,374414564 | 2,670836276 | Downregulated |
| 0,000158766 | 0,010805393 | -1,416296798 | 0,374672813 | 2,668995361 | Downregulated |
| 0,000297071 | 0,016766382 | -1,411518952 | 0,375915693 | 2,660170934 | Downregulated |
| 2,3847E-05 | 0,002691808 | -1,411205753 | 0,37599731 | 2,659593493 | Downregulated |
| 0,001492534 | 0,049621495 | -1,40901644 | 0,376568325 | 2,655560578 | Downregulated |
| 1,01275E-05 | 0,001358548 | -1,406279021 | 0,377283517 | 2,650526604 | Downregulated |
| 0,001361105 | 0,047541094 | -1,404235477 | 0,377818309 | 2,646774852 | Downregulated |
| 0,000397099 | 0,020195305 | -1,399918375 | 0,378950581 | 2,638866516 | Downregulated |
| 0,000345631 | 0,017972809 | -1,399060563 | 0,379175969 | 2,637297937 | Downregulated |
| 0,000349538 | 0,018074428 | -1,397929126 | 0,379473455 | 2,635230441 | Downregulated |
| 0,000110267 | 0,008723307 | -1,39631627 | 0,379897923 | 2,63228604 | Downregulated |
| 0,001242785 | 0,045288261 | -1,384008303 | 0,383152785 | 2,609924916 | Downregulated |
| 1,1358E-05 | 0,00150185 | -1,37421498 | 0,385762556 | 2,592268183 | Downregulated |
| 0,000538107 | 0,024903573 | -1,37405724 | 0,385804736 | 2,591984768 | Downregulated |
| 0,00064981 | 0,028237762 | -1,374035894 | 0,385810445 | 2,591946418 | Downregulated |
| 0,000157039 | 0,010793828 | -1,365423676 | 0,388120441 | 2,576519795 | Downregulated |
| 0,000865665 | 0,035297761 | -1,344933643 | 0,393672095 | 2,540185126 | Downregulated |
| 0,000306467 | 0,016875017 | -1,339290974 | 0,39521484 | 2,530269357 | Downregulated |
| 3,93453E-05 | 0,003951512 | -1,330623337 | 0,397596418 | 2,515113204 | Downregulated |

| | | | | | |
|---|---|---|---|---|---|
| 4,01653E-06 | 0,000688463 | -1,3286237 | 0,398147885 | 2,511629565 | Downregulated |
| 0,000996187 | 0,039404715 | -1,328497861 | 0,398182615 | 2,511410499 | Downregulated |
| 1,39887E-06 | 0,000275487 | -1,317648488 | 0,401188321 | 2,492594995 | Downregulated |
| 0,000345409 | 0,017972809 | -1,31291283 | 0,402507388 | 2,484426445 | Downregulated |
| 0,000305746 | 0,016875017 | -1,305614756 | 0,404548687 | 2,471890364 | Downregulated |
| 0,001089034 | 0,042018068 | -1,303316029 | 0,405193791 | 2,467954897 | Downregulated |
| 0,001037358 | 0,040513847 | -1,294559354 | 0,40766066 | 2,453020607 | Downregulated |
| 6,45799E-06 | 0,000964115 | -1,288261405 | 0,409444154 | 2,442335516 | Downregulated |
| 0,000181639 | 0,012182963 | -1,283204516 | 0,410881843 | 2,433789707 | Downregulated |
| 0,000442965 | 0,021579371 | -1,272636203 | 0,413902767 | 2,416026373 | Downregulated |
| 4,19578E-05 | 0,004088014 | -1,271133752 | 0,414334038 | 2,413511585 | Downregulated |
| 0,000128638 | 0,009589296 | -1,269275952 | 0,414867931 | 2,410405637 | Downregulated |
| 0,000561807 | 0,025616196 | -1,267091005 | 0,41549672 | 2,406757865 | Downregulated |
| 2,81212E-07 | 7,88758E-05 | -1,263465364 | 0,41654222 | 2,400717029 | Downregulated |
| 0,000432031 | 0,021158081 | -1,262972778 | 0,416684466 | 2,39989748 | Downregulated |
| 0,000129501 | 0,009589296 | -1,246599866 | 0,421440286 | 2,372815399 | Downregulated |
| 0,000109386 | 0,008723307 | -1,239532195 | 0,42350996 | 2,361219554 | Downregulated |
| 2,22665E-05 | 0,002544423 | -1,234558677 | 0,42497248 | 2,353093548 | Downregulated |
| 0,000242696 | 0,014575418 | -1,227848429 | 0,426953711 | 2,342174279 | Downregulated |
| 0,000339052 | 0,01793292 | -1,208518895 | 0,432712621 | 2,31100262 | Downregulated |
| 0,000803636 | 0,033356298 | -1,204215786 | 0,434005194 | 2,30411989 | Downregulated |
| 8,31095E-06 | 0,001165548 | -1,202404972 | 0,434550282 | 2,301229663 | Downregulated |
| 4,36906E-05 | 0,004169071 | -1,19963017 | 0,435386877 | 2,296807856 | Downregulated |
| 0,000135797 | 0,00981983 | -1,191948314 | 0,437711346 | 2,284610643 | Downregulated |
| 1,16327E-06 | 0,000239272 | -1,187157839 | 0,439167184 | 2,277037166 | Downregulated |
| 0,000604042 | 0,026879868 | -1,171242384 | 0,44403879 | 2,252055504 | Downregulated |
| 0,000680843 | 0,029040953 | -1,148984763 | 0,450942452 | 2,217577866 | Downregulated |
| 1,82414E-05 | 0,002192754 | -1,145665352 | 0,451981193 | 2,212481439 | Downregulated |
| 0,00023101 | 0,014380655 | -1,142587737 | 0,452946406 | 2,207766716 | Downregulated |
| 4,25768E-05 | 0,004105109 | -1,139183662 | 0,454016407 | 2,202563576 | Downregulated |
| 3,9703E-05 | 0,003951512 | -1,132625442 | 0,456084978 | 2,192573859 | Downregulated |
| 0,000902595 | 0,036323565 | -1,123675291 | 0,45892322 | 2,179013735 | Downregulated |
| 0,001007433 | 0,039680002 | -1,111025666 | 0,462964775 | 2,159991547 | Downregulated |
| 0,000873885 | 0,035393593 | -1,109070788 | 0,463592526 | 2,157066701 | Downregulated |
| 0,000829904 | 0,03414041 | -1,106990886 | 0,464261359 | 2,153959146 | Downregulated |
| 0,000461565 | 0,022251275 | -1,096820548 | 0,467545752 | 2,138828118 | Downregulated |
| 1,1539E-05 | 0,001504294 | -1,096646183 | 0,467602263 | 2,138569633 | Downregulated |
| 0,001403212 | 0,048104193 | -1,096380995 | 0,467688223 | 2,13817657 | Downregulated |
| 0,001438261 | 0,048585911 | -1,08107023 | 0,472678048 | 2,115604911 | Downregulated |
| 1,8045E-05 | 0,002192754 | -1,076442076 | 0,47419683 | 2,108828943 | Downregulated |
| 0,001127376 | 0,042525682 | -1,071091795 | 0,475958669 | 2,101022766 | Downregulated |
| 0,000157429 | 0,010793828 | -1,068419335 | 0,476841156 | 2,097134418 | Downregulated |
| 0,000301975 | 0,016875017 | -1,060201292 | 0,479565144 | 2,085222442 | Downregulated |
| 0,000689103 | 0,029124848 | -1,054464318 | 0,481475964 | 2,076946879 | Downregulated |
| 0,000215552 | 0,013855228 | -1,049011016 | 0,483299358 | 2,069110962 | Downregulated |
| 2,68883E-05 | 0,002962837 | -1,04256506 | 0,485463569 | 2,059886804 | Downregulated |
| 0,00013903 | 0,009888229 | -1,032289921 | 0,488933472 | 2,045268032 | Downregulated |
| 0,000939978 | 0,037664226 | -1,032065192 | 0,489009639 | 2,044949466 | Downregulated |
| 6,68749E-05 | 0,006009647 | -1,027553672 | 0,490541238 | 2,038564596 | Downregulated |
| 0,000638838 | 0,027891905 | -1,026444185 | 0,490918628 | 2,036997465 | Downregulated |

| | | | | | |
|---|---|---|---|---|---|
| 0,00074187 | 0,031071278 | -1,014659453 | 0,494945146 | 2,020425916 | Downregulated |
| 0,000328273 | 0,017462633 | -1,011237713 | 0,496120435 | 2,01563961 | Downregulated |
| 2,58503E-05 | 0,002882774 | -1,010598572 | 0,496340274 | 2,014746843 | Downregulated |
| 0,000230519 | 0,014380655 | -1,007369367 | 0,497452484 | 2,010242248 | Downregulated |
| 0,000875663 | 0,035393593 | -1,002187195 | 0,49924255 | 2,003034396 | Downregulated |
| 2,75701E-05 | 0,003002224 | -0,995353308 | 0,501613017 | 1,993568679 | Downregulated |
| 0,001197004 | 0,043966163 | -0,993710311 | 0,502184599 | 1,991299619 | Downregulated |
| 0,001160399 | 0,043121267 | -0,98518304 | 0,505161627 | 1,979564452 | Downregulated |
| 0,001368897 | 0,047633507 | -0,982197654 | 0,506208047 | 1,975472351 | Downregulated |
| 0,000123429 | 0,009441836 | -0,970168478 | 0,510446449 | 1,959069362 | Downregulated |
| 0,001338811 | 0,047479073 | -0,967952643 | 0,511231046 | 1,956062738 | Downregulated |
| 0,000191271 | 0,01246763 | -0,957233976 | 0,51504344 | 1,941583799 | Downregulated |
| 5,08902E-05 | 0,00475798 | -0,955132736 | 0,515794132 | 1,938758002 | Downregulated |
| 0,000380885 | 0,019585937 | -0,952912666 | 0,516588465 | 1,935776868 | Downregulated |
| 0,001355432 | 0,047522268 | -0,935523528 | 0,522852701 | 1,912584552 | Downregulated |
| 0,001193969 | 0,043966163 | -0,925270994 | 0,526581596 | 1,899040922 | Downregulated |
| 0,000327017 | 0,017462633 | -0,917497416 | 0,529426599 | 1,888835963 | Downregulated |
| 0,001115679 | 0,042525682 | -0,914410162 | 0,530560743 | 1,884798325 | Downregulated |
| 0,000970182 | 0,038633903 | -0,887698098 | 0,540475791 | 1,850221633 | Downregulated |
| 0,000289644 | 0,016549068 | -0,886755029 | 0,540829207 | 1,849012566 | Downregulated |
| 5,54462E-05 | 0,005132096 | -0,872709621 | 0,546120182 | 1,83109878 | Downregulated |
| 0,000542492 | 0,024981604 | -0,868431705 | 0,547741955 | 1,825677203 | Downregulated |
| 0,000587162 | 0,026511077 | -0,846367723 | 0,556183279 | 1,797968472 | Downregulated |
| 0,000611181 | 0,027067404 | -0,832060169 | 0,561726524 | 1,780225711 | Downregulated |
| 0,000497062 | 0,023236371 | -0,828939178 | 0,562943025 | 1,7763787 | Downregulated |
| 0,001265425 | 0,045753014 | -0,805670392 | 0,572096177 | 1,747957845 | Downregulated |
| 9,86019E-05 | 0,008076631 | -0,80418426 | 0,572685801 | 1,746158186 | Downregulated |
| 0,000268422 | 0,0156259 | -0,77873086 | 0,582879327 | 1,715620976 | Downregulated |
| 0,000582162 | 0,026414155 | -0,76982116 | 0,586490173 | 1,705058407 | Downregulated |
| 0,001424813 | 0,048307942 | -0,710158163 | 0,611253123 | 1,63598346 | Downregulated |
| 0,001298535 | 0,046406343 | -0,696401037 | 0,617109737 | 1,620457336 | Downregulated |
| 0,001344249 | 0,047489968 | -0,693423791 | 0,618384562 | 1,617116695 | Downregulated |

| ***Gene name*** |
|---|
| ccl27a |
| BX649355.1 |
| abcc8b |
| si:dkey-102c8.3 |
| CR382325.1 |
| sst1.1 |
| BX000363.1 |
| CR394546.4 |
| rgs5b |
| st6galnac1.2 |
| msrb1a |
| si:dkey-238k10.2 |
| BX248511.1 |
| CU457819.3 |
| calb1 |
| her8.2 |
| cxcl18b |
| f10 |
| flnca |
| bzw1a |
| si:dkey-25o1.7 |
| zmp:0000000634 |
| ccdc83 |
| CR354588.1 |
| mt-atp8 |
| prozb |
| TCIM |
| znf644a |
| gpx4a |
| ncanb |
| adgb |
| stat3 |
| her13 |
| si:ch73-138i16.1 |
| pdk2b |
| si:ch1073-190k2.1 |
| pcgf1 |
| ccndx |
| kidins220a |
| grm6b |
| her4.1 |
| cox5b2 |
| sun1 |
| cyp46a1.2 |
| cryabb |
| ier3ip1 |
| ntrk3a |
| ucn3l |
| BX324206.3 |

Upregulated in mutant

downregulated in muta

| |
|---|
| pclob |
| taco1 |
| cox4i1l |
| gcnt4a |
| si:ch211-160o17.4 |
| hist1h2a11 |
| rab7a |
| zgc:171699 |
| pvalb9 |
| AL808129.1 |
| s100a1 |
| si:dkey-174n20.1 |
| dusp2 |
| top1l |
| asmtl |
| ppargc1b |
| CR769769.2 |
| snf8 |
| lmo7a |
| fstl1b |
| sall3a |
| morn5 |
| mt-co2 |
| sumo3b |
| zgc:194246 |
| stmn1a |
| si:ch73-350k19.1 |
| si:dkeyp-67f1.1 |
| abhd12 |
| irf7 |
| mt-nd3 |
| entr1 |
| as3mt |
| hipk3a |
| mt-nd1 |
| pbld2 |
| zgc:112234 |
| sall1a |
| six2b |
| ndufa1 |
| polr2l |
| lyrm7 |
| si:ch211-145b13.5 |
| lsm5 |
| CU467110.1 |
| xpo1b |
| CU694532.1 |
| pde4ba |
| cox7c |
| mt-atp6 |

| |
|---|
| mt-nd5 |
| cabp5b |
| zgc:112234 |
| fam32a |
| rbfa |
| acbd7 |
| cops9 |
| si:dkey-27p23.3 |
| mt-co1 |
| atf3 |
| eif1b |
| tmem14ca |
| mt-cyb |
| ppp1r15a |
| mvda |
| C11H3orf14 |
| chrac1 |
| CU459186.2 |
| ssx2ipb |
| CR536604.1 |
| mt-nd2 |
| BX530018.2 |
| pllp |
| cdk5rap1 |
| f11r.1 |
| si:ch211-274p24.2 |
| gpx2 |
| *micos13* |
| *coa6* |
| *C5H11orf1* |
| *cox6c* |
| *cox7b* |
| s100s |
| *mt-co3* |
| *CR383676.2* |
| *mt-nd4* |
| vamp8 |
| pou4f2 |
| pax6a |
| itgb1b.2 |
| si:ch211-222e20.4 |
| rbpms2a |
| irx4a |
| isl2b |
| mab21l1 |
| tubb5 |
| cldne |
| ttc23 |
| gng13b |
| si:dkey-172m14.1 |

| |
|---|
| dtwd2 |
| cspg5a |
| celf3b |
| gnao1b |
| nup58 |
| st13 |
| pax10 |
| fbxw2 |
| ppp1r14ba |
| snap25b |
| krt4 |
| dtnbp1b |
| marcksl1a |
| sgce |
| calb2a |
| foxg1b |
| islr2 |
| hist1h4l |
| fam149a |
| hmx4 |
| atg7 |
| si:dkey-23a13.22 |
| FNDC10 |
| si:dkey-56f14.7 |
| gnai2b |
| stmn2b |
| gsdmeb |
| fgf3 |
| tbx1 |
| pjvk |
| cttn |
| fam53b |
| pcp4a |
| pggt1b |
| kcmf1 |
| foxg1c |
| cep19 |
| mrm3a |
| vps4a |
| grem2b |
| ccdc15 |
| npm1b |
| mycn |
| chmp1a |
| herpud1 |
| tbc1d7 |
| elavl3 |
| samm50l |
| icn |
| lfng |

| |
|---|
| spry2 |
| PRRC2B |
| mb |
| bysl |
| ubl7a |
| zgc:101562 |
| twf1a |
| rbpms2b |
| cox15 |
| ankrd46b |
| bmp5 |
| tp53inp1 |
| ildr1a |
| tectb |
| tspan7b |
| march4l |
| ctnnbip1 |
| sox21a |
| mybphb |
| acads |
| znf503 |
| grtp1a |
| alkbh4 |
| alas1 |
| marcksl1b |
| si:ch211-151p13.8 |
| scamp2 |
| isl1 |
| galnt18b |
| stk3 |
| zgc:158291 |
| npepl1 |
| pias2 |
| PSMD2 |
| barhl1b |
| stard10 |
| sox2 |
| gpm6ab |
| si:ch211-284e13.6 |
| si:ch211-212o1.2 |
| rnf170 |
| azin1b |
| fgf13b |
| nol12 |
| psmd3 |
| drap1 |
| NA |
| cdkn1ba |
| pkma |
| zgc:65894 |

| |
|---|
| nanos1 |
| pcbp2 |
| zgc:112234 |
| tmeff1b |
| chtopa |
| rtn1b |
| ethe1 |
| zdhhc13 |
| klf5l |
| cdo1 |
| march5l |
| atoh1a |
| capzb |
| gadd45gb.1 |
| sbf1 |
| nars1 |
| sox1a |
| shisa2a |
| ift74 |
| cdcp2 |
| TUBB4B |
| creb3l3l |
| odc1 |
| BX323060.3 |
| bdnf |
| slc25a39 |
| tuba1a |
| si:ch211-243g18.2 |
| elmo1 |
| si:ch211-114n24.6 |
| sucla2 |
| cct4 |

| P-adj | Abs(FC) | Gene name | Function |
|---|---|---|---|
| 0,009888229 | 2,733792803 | **gsdmeb** | **Involved in inner ear morphogenesis** and regulation of biosynthetic process. Is expressed in heart tube; intermediate cell mass of mesoderm; nervous system; neural crest; and **otic vesicle.** Human ortholog(s) of this gene **implicated in autosomal dominant nonsyndromic deafness 5 and sensorineural hearing loss**. Orthologous to human GSDME (gasdermin E). |
| 0,000219095 | 2,714264623 | **pjvk** | Human ortholog(s) of this gene implicated in autosomal recessive nonsyndromic deafness 59. Orthologous to human PJVK (pejvakin). |
| 0,025616196 | 2,406757865 | **ildr1a** | Predicted to localize to integral component of membrane. Is expressed in EVL; ectoderm; and sensory system. Human ortholog(s) of this gene implicated in autosomal recessive nonsyndromic deafness 42. Orthologous to human ILDR1 |

<table>
<tr><th>Expression</th><th></th></tr>
<tr><td>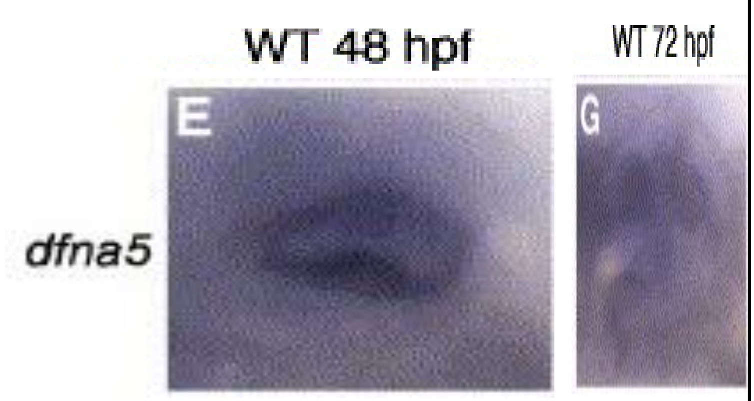
</td><td>expression in zebrafish</td></tr>
<tr><td>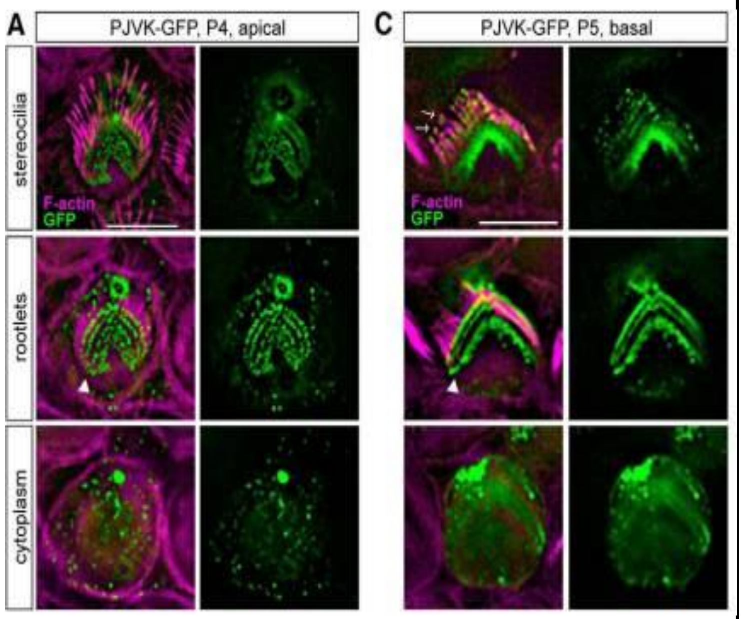
</td><td>expression in mice</td></tr>
<tr><td>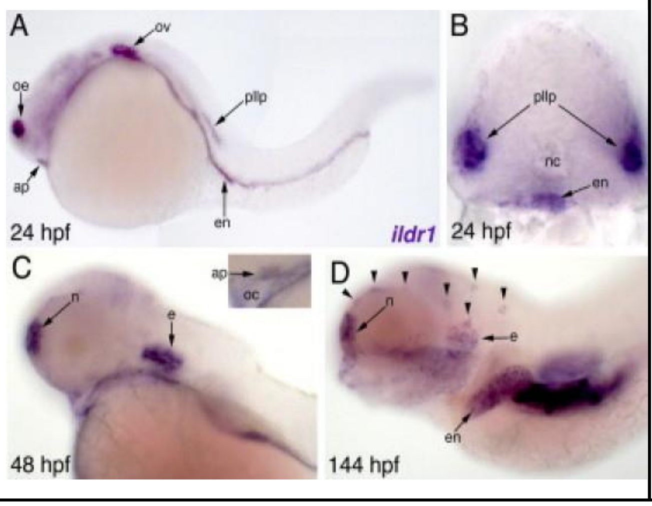
</td><td>expression in zebrafish</td></tr>
</table>

<table>
<tr><td>pore formation and involvement in pyropoptosis (inflammatory apoptosis)</td><td rowspan="2">Same family (Gasdermin family)</td></tr>
<tr><td>Recent studies have ascribed a role for pejvakin in the oxidative-stress induced proliferation of peroxisomes in hair cells and auditory neurons in response to noise exposure. Pejvakin selectively localizes to stereociliary rootlets and is required to preserve the integrity of mechanosensitive stereocilia</td></tr>
<tr><td>ILDR1 is playing a major role in the development of auditory hair cells, semicircular canal and tricellular tight junction. It has been reported that IDLR1 protein may involve as barrier for cellular tight junction (Higashi et al., 2015). In addition, Kim et al. reported that mutation in ILDR1 causes disruption of tricellulin, hence, the formation of tricellular tight junction can’t be possible (Kim et al., 2015).</td><td></td></tr>
</table>
